# Cell surface ribonucleoproteins cluster with heparan sulfate to regulate growth factor signaling

**DOI:** 10.1101/2024.07.25.605163

**Authors:** Peiyuan Chai, Jonathan Perr, Lauren Kageler, Charlotta G. Lebedenko, Joao M.L. Dias, Eliza Yankova, Jeffrey D. Esko, Konstantinos Tzelepis, Ryan A. Flynn

## Abstract

Receptor-ligand interactions govern a wide array of biological pathways, facilitating a cell’s ability to interrogate and integrate information from the extracellular space. Here, using an unbiased genome-wide knockout screen, we identify heparan sulfate proteoglycans (HSPGs) as a major component in the organizational mechanism of cell surface glycoRNA and cell surface RNA binding proteins (csRBPs). Cleavage of mature heparan sulfate chains, knockout of *N-* and 6-*O*-sulfotransferases, overexpression of endo-6-*O*-sulfatases, or the addition of exogenous heparan sulfate chains with high 2-*O* sulfation result in marked loss in glycoRNA-csRBP clustering in U2OS cells. Functionally, we provide evidence that signal transduction by HS-dependent growth factors such as VEGF-A_165_ is regulated by cell surface RNAs, and in vitro VEGF-A_165_, selectively interacts with glycoRNAs. Our findings uncover a new molecular mechanism of controlling signal transduction of specific growth factors across the plasma membrane by the regulated assembly of glycoRNAs, csRBPs, and heparan sulfate clusters.

## Introduction

Cells leverage biophysical and regulatory interfaces to communicate with the extracellular environment. Heparan sulfate proteoglycans (HSPGs), a key class of cell surface glycoconjugates, consist of a core protein and one or more covalently linked heparan sulfate (HS) chains^1^. Most of the HSPG core proteins are membrane-associated, anchored by way of a transmembrane domain (e.g., syndecans) or by a glycosylphosphatidylinositol (GPI) anchor (e.g. glypicans)^2^. The protein interactome of HSPGs is well established and includes chemokines, cytokines, growth factors^3,4^, morphogens^5,6^, cell adhesion proteins^7–9^, heat shock proteins^10^, and viral components like the SARS-CoV-2 spike protein^11^. HSPGs are thought of conceptually as coreceptors that facilitate the formation of ligand-receptor complexes. A major source of control over protein-HS interactions occurs via sulfated domains of the chains, affected by the total extents of sulfation as well as the arrangement of sulfated residues in the chains^6^. HS and HSPGs are critical to a diverse set of biological processes, spanning development, physiology, and pathophysiology.

While the expression of the protein component of HSPGs can control some aspects of HSPG biology, the particularities of the carbohydrate polymer have been found to be critical for many of the functional aspects of HSPGs. HS chains vary enormously in terms of length and degree of sulfation. Its assembly occurs in the Golgi apparatus in a template-independent manner. A series of enzymes initiate HS chain formation, eventually leading to a core glucuronic acid-galactose-xylose tetrasaccharide linkage region^12^. EXTL3 initiates the formation of the repeating disaccharide units of HS by transfer of the first N-acetyl-D-glucosamine (GlcNAc) unit to the linkage region tetrasaccharide. A heterodimer complex of EXT1 and EXT2 adds alternating residues of D-glucuronic acid (GlcA) and GlcNAc to the nascent polymer^13–15^. Sulfation of the chains is initiated by the one or more members of the NDST family of N-deacetylases-N-sulfotransferases acting on a subset of GlcNAc residues, followed by partial epimerization of adjacent GlcA residues to l-Iduronic acid (IdoA) followed by 2-*O*-sulfation (catalyzed by Hs2st), 6-*O*-sulfation of GlcNAc and GlcNS residues (by Hs6st1-3), and occasional 3-*O*-sulfation of GlcNS residues (by Hs3st1, 2, 3a, 3b, 4, 5, or 6)^12,16^.

High levels and/or site-specific sulfation of HS drive the various activities ascribed to HSPGs^6^. We recently described a new negatively charged glycopolymer on the cell surface called sialoglycoRNA^17^, which present sialylated and fucosylated N-glycans on small RNAs, covalently attached to one another via the modified RNA base 3-(3-amino-3-carboxypropyl)uridine (acp3U)^18^. On the cell surface, these sialoglycoRNAs are in physical proximity to specific RNA binding proteins on the plasma membrane of cells (csRBPs)^19^. The cell penetrating peptide (CPPs) TAT localizes to and enters cells in a manner partially dependent on RNA at sites where csRBPs cluster^19^. CPPs are classically reported to leverage HSPGs for cell surface association and entry^20^, highlighting the possibility that glycoRNA-csRBP clusters and HSPGs have some relationship. More broadly, HSPGs are well characterized to bind extracellular growth factors leading to subsequent signal transduction by forming a growth factor-growth factor receptor complex on the cell surface^6^. Among these HS-binding growth factors, specific proteoforms of vascular endothelial growth factor (VEGF) promotes angiogenesis, endothelial cell migration, and proliferation; VEGF binding depends on 6-*O*-sulfation of HS^21,22^. VEGF receptors (VEGFR) also occur as clusters on the cell surface^23,24^, which may influence VEGF-VEGFR activation of Ras/Raf1/MEK, which in turn phosphorylates ERK1/2^3^. Activated ERK has many effects on cells, including regulating cell growth, inflammatory signals, and cell death^25^, however is it not clear if or how cell surface RNAs may impact these pathways.

Here we take a genetic approach to dissect the identity and assembly of glycoRNA-csRBP clusters. We define tools that enable examination of these clusters on live cells and apply them to perform a genome-wide knockout screen. Our screening effort revealed a major genetic dependency of glycoRNA-csRBP clusters on heparan sulfate biogenesis, a mechanism conserved across multiple cell types. Selective and rapid cleavage of heparan sulfate chains with heparin lyase diminished glycoRNA-csRBP clusters. Colocalization and time course experiments suggest HSPGs are sites of assembly for glycoRNA-csRBP clusters. Interrogation of the specific types of sulfation revealed that 6-*O*-sulfation is key to promoting the formation of glycoRNA-csRBP clusters. Finally, we show that the growth factor VEGF-A_165_ can directly bind RNA on the cell surface and can interact with glycoRNAs *in vitro*. VEGF-A_165_ signaling, which traditionally is thought to leverage HSPGs for cell surface interaction, can be modulated glycoRNAs.

## Results

### Siglec-11 binds cells in an RNA-dependent manner and is in proximity to glycoRNAs on the cell surface

We previously found that Siglec-11 (recombinant form of the extracellular domain of Siglec-11 fused to a human Fc domain) binds to the surface of HeLa cells in an RNA-dependent manner^17^. To expand our understanding of which Siglecs have RNA-dependent cell surface binding, we screened the 13 commercially available Siglec-Fc fusion proteins for binding to suspension (MOLM-13) and adherent (U2OS) cell lines. Siglec-4, Siglec-7, Siglec-9, and Siglec-11 strongly bound both cell types above the level shown by the control IgG-Fc (**Figure 1A, S1A**). A pooled RNase treatment^19^ (RNase A, a single stranded RNase and RNase III, a double stranded RNase) resulted in a significant reduction of Siglec-11 binding ability, as measured by total dots per cell (**Figure 1A**, **1B**) and intensity per cell (**Figure 1A, S1B**), on both cell types. Live cell RNase treatment had no impact on the binding of Siglec-4, Siglec-7, or Siglec-9 (**Figure 1A**, **1B, S1B**). These data support the initial observation of RNA-dependent binding of Siglec-11. As a control for reagent specificity, we also treated live cells with a sialidase cocktail (**Methods**) to evaluate the impact of removal of cell surface sialic acids. Both Siglec-7 binding and cell surface periodate labeling^26^ demonstrated robust and significant loss of binding after sialidase treatment (**Figure S1C**), whereas Siglec-11 binding was not impacted (MOLM-13) or only mildly impacted (U2OS, **Figure S1C**) under the same conditions.

**Figure 1.**
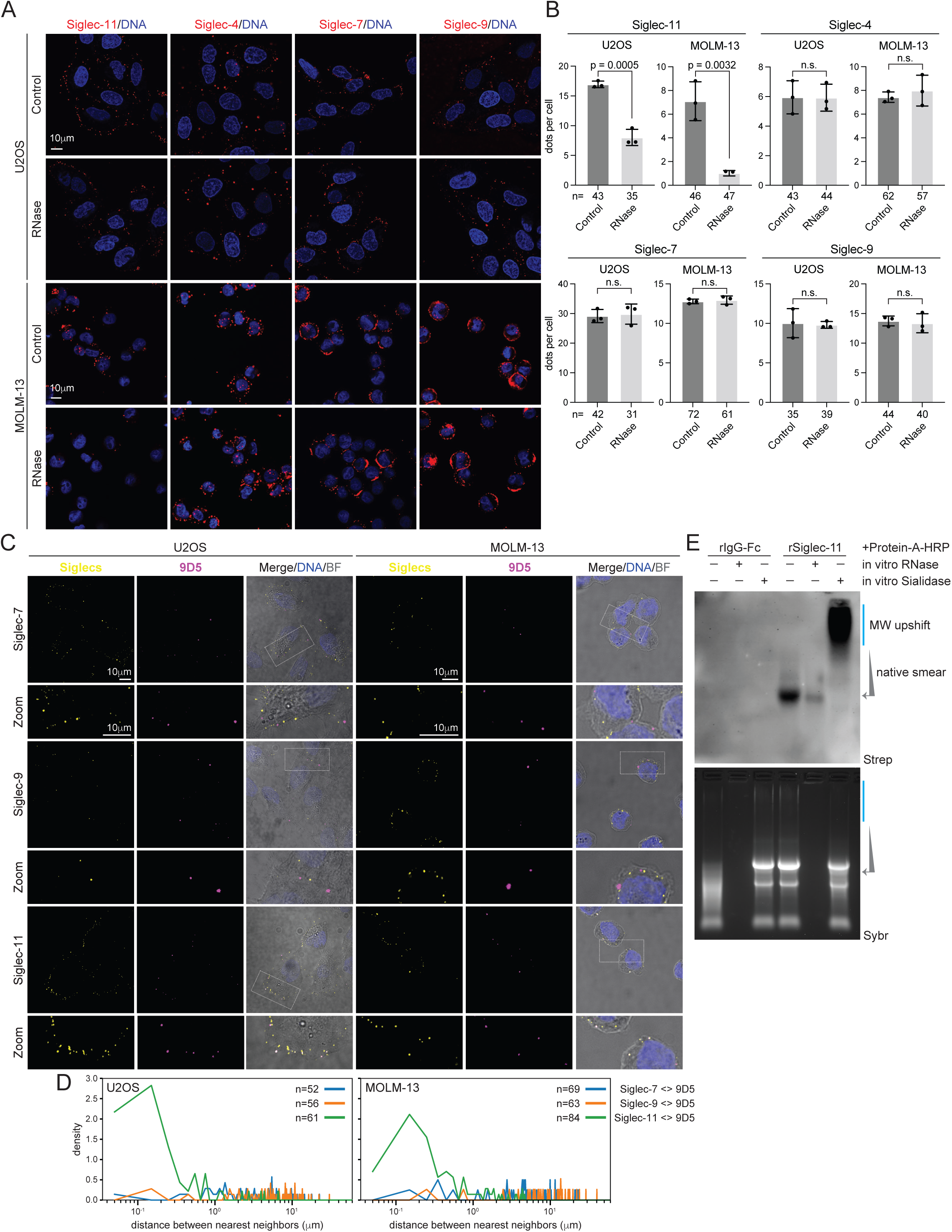
Siglec-11 is a major RNA binding Siglec on the cell surface. A. Representative confocal images of U2OS and MOLM-13 cells treated with RNase A and RNase III (RNase pool) for 30 min and stained live with the indicated Siglec-Fc reagents (red). DNA was stained with DAPI (blue). Scale bar, 10 µm. B. Quantification of the indicated Siglec dot numbers in U2OS and MOLM-13 cells treated with RNase A and RNase III for 30 min from 3 independent experiments with the number (n) of cells analyzed noted. C. Representative confocal images of U2OS and MOLM-13 cells co-stained with the indicated Siglec-Fc reagents (yellow) and 9D5 (magenta). DNA was stained with DAPI (blue). An enlargement of the hatched box is shown, and a bright field (BF) is shown to represent the outline of the cell membrane. Scale bar, 10 µm. D. Nearest neighbor distance analysis of the Siglec pairs imaged in (C). For each pair, the distance (nanometers, nm) from that anchor (left side protein name in the figure key) to the other pair was calculated across the indicated number of cells. These values were plotted in a density histogram. E. RNA blotting of MOLM-13 cells stained while alive with IgG-Fc or Siglec-11 and labeled with biotin aniline to tag cell surface RNAs. Whole cell RNA was then extracted, processed in vitro with RNase or sialidase, and then analyzed by gel. Total RNA (bottom) and biotin (top) images highlight selective labeling of glycoRNA material only with Siglec-11.

Examining the confocal imaging data of Siglec-11 binding showed the ligands of Siglec-11 formed as clusters or puncta on the cell surface (**Figure 1A**). Cell surface puncta of Siglec ligands has been seen previously, for example with super resolution imaging of Siglec-7 and Siglec-9 binding^27^. Given the observed punctate nature of Siglec-11 binding and its RNA-dependency, we predicted that RNA should colocalize with Siglec-11 on the cell surface. We recently described a new domain on the cell surface, glycoRNA-csRBP clusters^19^, where cell surface RNA binding proteins (csRBPs) and glycoRNAs colocalize to facilitate the functional entry of molecules such as cell penetrating peptides. To address the possible association of Siglec-11 ligands near or within these glycoRNA-csRBP clusters, we used an anti-dsRNA antibody (9D5) that detects cell surface RNA^19^. Costaining of U2OS and MOLM-13 cells with 9D5 and Siglec-11 showed that 42% and 61% of 9D5 puncta overlapped with Siglec-11 on MOLM-13 and U2OS cells, respectively. Co-staining 9D5 with Siglec-7 and separately with Siglec-9, both of which are not sensitive to RNases (**Figure 1A**, **1B, S1B**) showed that only 2.5% and 1.4% of 9D5 puncta overlapped with Siglec-7 or Siglec-9 in MOLM-13, and 0% and 2.8% of 9D5 puncta overlapped with Siglec-7 or Siglec-9 in U2OS, respectively (**Figure 1C**, **1D**). The well-correlated binding of 9D5 and Siglec-11 suggests that Siglec-11 ligands are near RNA on the cell surface.

To directly assess if Siglec-11 ligands are in proximity to glycoRNAs, we performed cell surface proximity labeling, using biotin aniline to label RNAs in proximity to bound Siglec-11. Analysis of biotin signal from total RNA extracted from labeled cells showed that the IgG-Fc control had no signal, whereas Siglec-11 staining produced an extended smear to higher molecular weights from both MOLM-13 and U2OS cells (**Figure 1E, S1D**). *In vitro* digestion of total RNA from labeled cells demonstrated that the biotin signal was sensitive to RNase whereas sialidase treatment resulted in a more slowly migrating smear (**Figure 1E**). This RNase and sialidase sensitivity is consistent with previous results using plant lectins (WGA and MAA-II) to label cell surface RNAs^17^. Together these data demonstrate that Siglec-11 binding on living cells depends on cell surface RNA, and that the binding region is in proximity to glycoRNA, suggesting that Siglec-11 ligands are near to or within glycoRNA-csRBP clusters.

### Heparan sulfate biogenesis is a major genetic determinant of glycoRNA-csRBP clustering

To gain insight into the genetic basis of Siglec-11 and 9D5 binding, a genome-wide CRISPR-Cas9 gene knockout approach^28^ was developed based on flow cytometry of MOLM-13 cells using Siglec-11, 9D5, and the plant lectin MAA-I, which is well characterized to selectively bind sialic acid^29^. We analyzed sgRNA sequencing data from unsorted input and the bottom 5% of sorted cells and determined which sgRNAs and corresponding genes were enriched for reducing the binding of the three probes (**Figure 2A**, **2B, S2A**). Using a CRISPR-score cutoff of -0.8, we found 154, 187, and 246 hits in the Siglec-11, 9D5, and MAA-I screens, respectively (**Table S1**). The top hits enriched in the MAA-I screen were related to sialic acid biosynthesis (**Figure S2A**, **Table S1**), for example CMAS and NANS^25^, but both of these genes scored poorly in the Siglec-11 and 9D5 screen, confirming the specificity of the screen. Analysis of these enriched genes showed a more robust overlap between 9D5 and Siglec-11 enriched sgRNAs, compared to MAA-I with either of the other two probes (**Figure 2C**). Gene ontology analysis (GO) of the cellular compartment and biological process demonstrated that enriched genes were related to membrane compartments (**Figure S2B**), suggesting each probe was addressing cell surface biology. Inspection of the top hits of Siglec-11 revealed EXT1, EXT2, and UXS1 as the top three genes, which are key enzymes of heparan sulfate biogenesis (**Figure 2A**). The 9D5 screen also revealed EXT1 as a highly scoring hit, while UXS1 and EXT2 were present but below the -0.8 cutoff (**Figure 2B**).

**Figure 2.**
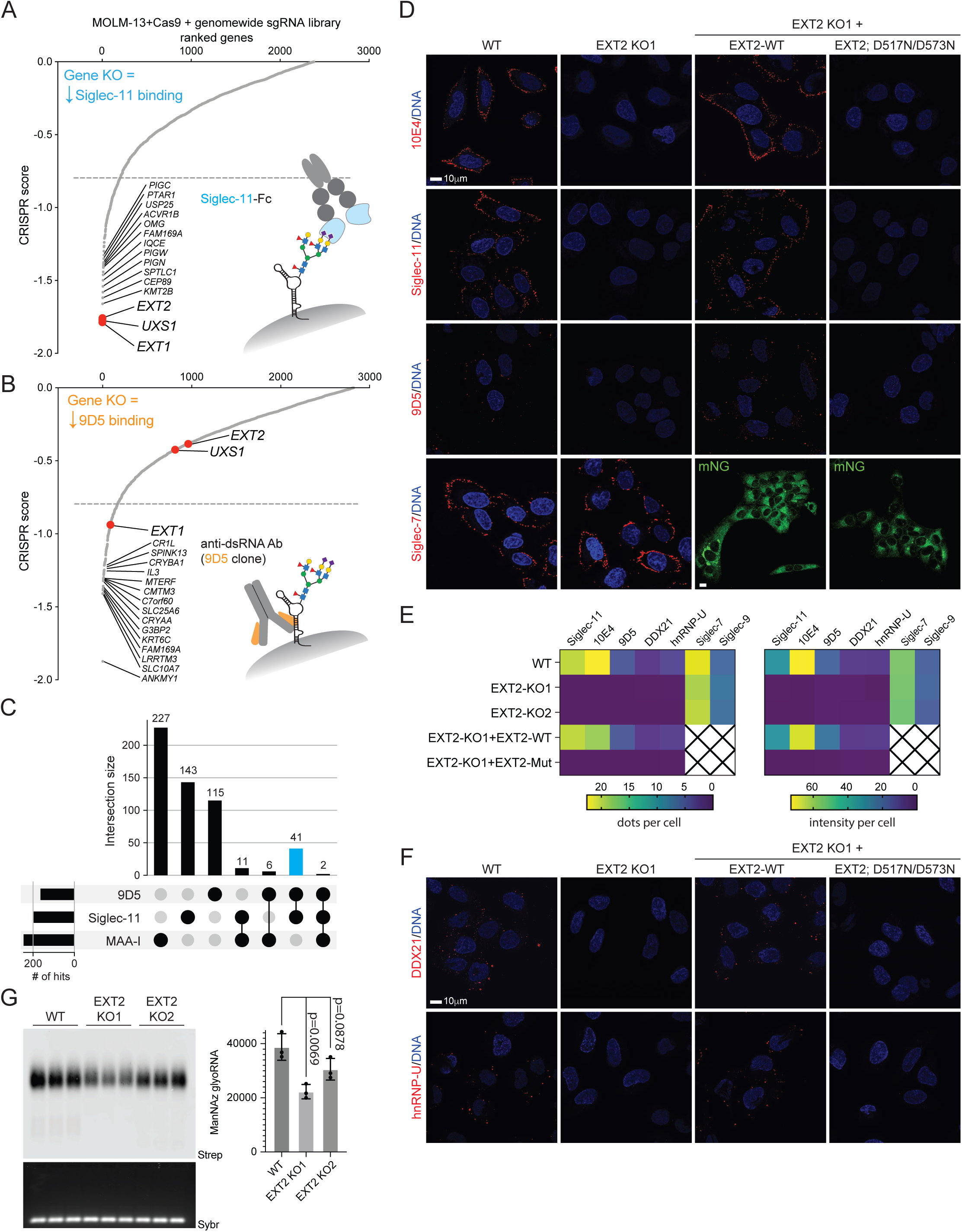
glycoRNA-csRBP clusters are dependent on heparan sulfate biogenesis. A. Dot plot of genes identified in the genome-wide knockout (KO) screen for loss of Siglec-11 cell surface binding ranked by CRISPR score. The top 15 gene names are displayed with a line drawn at the -0.8 score cut off. The inset cartoon illustrates how Siglec-11 could interact with a cell surface glycoRNA. B. Dot plot of genes identified in the genome-wide KO screen as in (A), here for the loss of 9D5 cell surface binding. The inset cartoon illustrates how 9D5 could interact with a cell surface glycoRNA. C. Upset plot analysis of genes with a score cutoff of -0.8 from the Siglec-11, 9D5, and MAA-I genome-wide KO screens. The common overlapping hits between 9D5 and Siglec-11 are highlighted in blue. The total number of hits for each intersection is noted. D. Representative confocal images of wild-type (WT), EXT2 knock-out (KO), EXT2 KO-mEmerald-EXT2, EXT2 KO-mEmerald-D517N/D573N EXT2 U2OS cells stained live with 10E4, Siglec-11 Fc, and 9D5, (all in red). DNA was stained with DAPI (blue). Scale bar, 10 µm. E. Quantification of 10E4, Siglec-11, 9D5, cs-DDX21, and cs-hnRNP-U dot numbers and intensity per cell from D and F from 3 independent experiments. F. Representative confocal images of WT, EXT2 KO, EXT2 KO-mEmerald-EXT2, EXT2 KO-mEmerald-D517N/D573N EXT2 U2OS cell lines stained live with anti-DDX21 and anti-hnRNP-U (both in red). DNA was stained with DAPI (blue). Scale bar, 10 µm. G. RNA blotting of U2OS small RNA labeled with Ac_4_ManNAz and detected with copper-free click of dibenzocyclooctyne-PEG4-biotin (DBCO-biotin). In gel detection of small RNA with SybrGold (Sybr, bottom) and on membrane detection of biotin (Strep, top) is shown. Quantification plotted on the right and statistical assessment of the intensities was performed with a t-test.

Both Siglec-11 and 9D5 demonstrated genetic dependency on heparan sulfate (HS) biosynthesis. The HS chain is initiated by xylosylation of proteoglycan core proteins, which depends on UDP-Xylose formation catalyzed by UXS1, and is subsequently elongated by a hetero-dimeric complex formed by EXT1 and EXT2 in the golgi apparatus^1^. We next generated two individual knockout clones of EXT2 in U2OS cells, to validate the effect of HS biogenesis on Siglec-11 and 9D5 binding outside of the genome-wide screen and in another cell type (U2OS, **Figure S2C, S2D, 2D**). We stained the live EXT2 knockout (KO) cells with 10E4, a specific antibody recognizing mature HS^30,31^, and verified that HS was not produced (**Figure 2D**, **2E, S2E, S2F**). Consistent with the high score for EXT2 in the genome-wide screen, EXT2 deficiency resulted in complete loss of both Siglec-11 and 9D5 binding (**Figure 2D**, **2E, S2E, S2F**). Binding of Siglec-7 and Siglec-9 was not affected (dots per cell or intensity per cell) in EXT2-KO cells (**Figure 2D**, **2E, S2E, S2F**). To assess if csRBPs were also regulated by HS biogenesis, we performed live cell staining using anti-DDX21 and anti-hnRNP-U antibodies. Consistent with 9D5 and Siglec-11 signals, no DDX21 and hnRNP-U antibody binding occurred on the cell surface in EXT2 KO cells (**Figure 2E**, **2F, S2F, S2G**). To understand if this effect was due to the enzymatic activity or some other role of EXT2, EXT knockout cells were transfected with WT EXT2 or a catalytically inactive (D517N/D573N^15^) mutant EXT2 cDNA. WT EXT2 rescued the loss of 9D5, Siglec-11, csDDX21, and cs-hnRNP-U signal (**Figure 2D-2F, S2D-S2G**), whereas the catalytically inactive mutant did not. This finding indicates that the N-acetylglucosaminyltransferase (GlcNAc-T) activity of EXT2 in HS polymerization is required to facilitate glycoRNA-csRBP clustering on the cell surface. Finally, we examined the levels of sialoglycoRNA in EXT2 KO cells. Loss of EXT2 led to partial (42.5% in KO1 and 21.2% in KO2) reduction in sialoglycoRNA signal (**Figure 2G**). Together these data suggest that HS biogenesis is required for glycoRNA-csRBP clustering on the cell surface and that there may be close physical interaction.

### glycoRNA-csRBP clusters are colocalized with, and dependent on, intact heparan sulfate polymers

Mature HS proteoglycans present on the cell surface or in the extracellular matrix after biogenesis and vesicular trafficking to the plasma membrane. To determine whether the mature HS chains structure on the cell surface or the intracellular HS biogenesis process facilitates glycoRNA-csRBP clustering on the cell surface, we performed live cell staining on U2OS cells after heparin lyase treatment (45 min, heparin lyases I/II/III), which cleaves heparan sulfate chains in the extracellular space. As predicted, binding of 10E4 was lost in heparin lyase-treated cells (**Figure 3A, S3A**). Heparin lyase treatment also completely removed the Siglec-11 and 9D5 binding on cells (**Figure 3A, S3A**), but did not affect Siglec-7 and Siglec-9 binding (**Figure 3A, S3A**). Staining with anti-DDX21 and anti-hnRNP-U antibodies showed that cleavage of mature HS chains also resulted in the loss of csDDX21 and cs-hnRNP-U puncta on the cell surface (**Figure 3A, S3A**). Given the robust sensitivity of 9D5 and Siglec-11 binding to heparin lyase, we tested the activity of the enzyme on glycoRNAs *in vitro*. Heparin lyase had no direct effect on glycoRNAs, indicating that HS was not covalently linked to the RNAs (**Figure S3B**). These data show that mature HS chains are required for glycoRNA-csRBP cluster formation.

**Figure 3.**
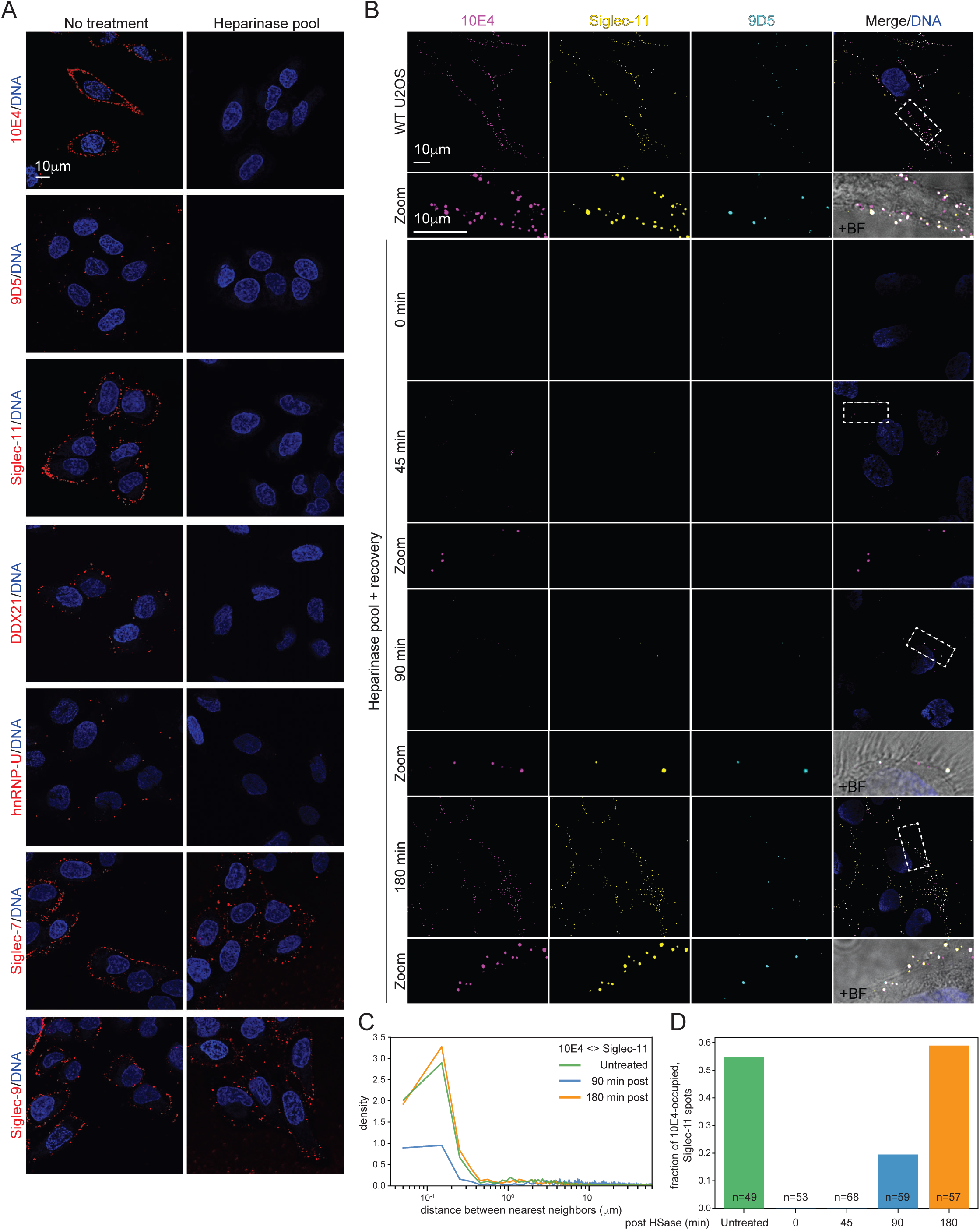
Cell surface glycoRNAs and csRBPs are dependent on intact heparan sulfate chains. A. Representative confocal images of U2OS cells treated with heparinase pool for 30 min, and stained live with 10E4, Siglec-11, 9D5, anti-DDX21, anti-hnRNP-U, Siglec-7-Fc, and Siglec-9-Fc (all in red). DNA was stained with DAPI (blue). Scale bar, 10 µm. B. Representative confocal images of U2OS cells treated with heparinase pool for 30 min, recovering for the indicated times, and co-stained with 10E4 (Cyan), Siglec-11 (yellow), and 9D5 (magenta). DNA was stained with DAPI (blue). An enlargement of the hatched box is shown, and a bright field (BF) is shown to represent the outline of the cell membrane. Scale bar, 10 µm. C. Nearest neighbor distance analysis of the 10E4 and Siglec-11 in (B). For each pair, the nm distance from 10E4 to Siglec-11 was calculated across. These values were plotted in a density histogram. D. Quantification fraction of 10E4-occupied, Siglec-11 dots in (B) from 3 independent experiments.

Given the co-localization of 9D5 and Siglec-11 and sensitivity to heparin lyases, we sought to understand the spatial relationship between 9D5, Siglec11, and HS. We therefore performed a three-color co-staining experiment in U2OS cells and found that Siglec-11 puncta are highly correlated in localization with 10E4 puncta (**Figure 3B**, **3C; green line**). We next assessed the temporal regulation on glycoRNA-csRBP clustering by conducting a recovery experiment in cells after heparin lyase treatment, removal of the enzymes and incubation of the cells with fresh media for 0, 45, 90, or 180 minutes. HS puncta reappeared up after a 45-minute recovery from heparinase treatment (9.4%) while Siglec-11 or 9D5 clustering was not detected (**Figure 3B**, **3C**, **3D**). After 90 minutes, 9D5 and Siglec-11 started to recover together at sites of large HS clusters. (**Figure 3B**, **3C**, **3D**). Finally, all the signals of HS, Siglec-11, and 9D5 recovered to normal after 180 minutes (**Figure 3C**, **3D**), confirming that cell surface HS chains are vital for the formation of glycoRNA-csRBP clusters.

### 6-*O*-sulfation of Heparan sulfate chains facilitates glycoRNA-csRBP cell surface clustering

Next, we examined how the sulfation of heparan sulfate chains participates in the glycoRNA-csRBP clusters. NDST1 catalyzes GlcNAc N-deacetylation/N-sulfation of the HS chains (**Figure 4A**). Deletion of NDST1 in U2OS cells (**Figure S4A, S4B**) resulted in a 74% loss of N-sulfo glucosamine residues as measured by reduction of 10E4 staining, which is known to depend on N-sulfation (**Figure 4B**, **4C**). Staining of NDST1 knockout cells with Siglec-11, 9D5, anti-DDX21, and anti-hnRNP-U revealed that NDST1 deficiency caused 77%, 72%, 62%, and 63% loss of dot number per cell in Siglec-11, 9D5, csDDX21, and cs-hnRNP-U (**Figure 4B**, **4C**). Because the NDST family has four members and U2OS cells express both NDST1 and NDST2^28^, we expected only a partial effect when NDST1 was deleted. To broadly inhibit sulfation, we treated cells with sodium chlorate, a metabolic inhibitor of sulfation^32^, and found near complete loss of 10E4, Siglec-11, 9D5, csDDX21, and cs-hnRNP-U, whereas sodium chloride (control) had no effect (**Figure S4C**).

**Figure 4.**
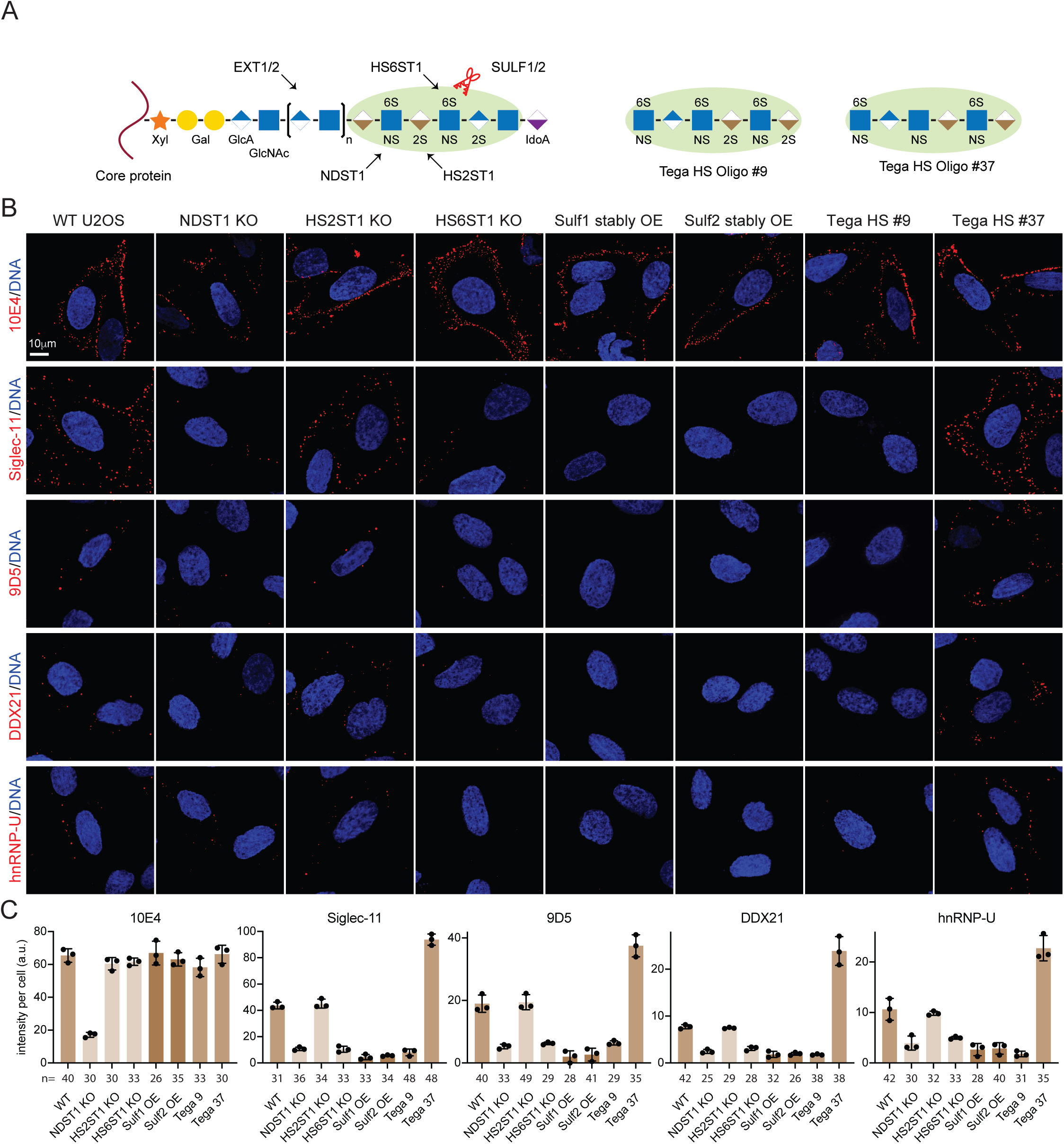
Proteoglycan sulfation modulates glycoRNA-csRBP clustering. A. Schematic of a heparan sulfate proteoglycan with the regions of activity for the various enzymes and HS oligos perturbed in the following experiments. B. Representative confocal images of WT, NDST1 KO, HS6ST1 KO, HS2ST1 KO, Sulf1 stably overexpressing (OE), Sulf2 stably overexpressing, and Tega HS #09 and Tega HS #37 treated U2OS cells stained with 10E4, Siglec-11, 9D5, anti-DDX21, and anti-hnRNP-U (all in red), separately. DNA was stained with DAPI (blue). Scale bar, 10 µm. C. Quantification of data in (B) with number of cells counted from 3 independent experiments.

To refine our understanding of the role of HS sulfation we next assessed how modulating uronyl 2-*O*-sulfation and glucosaminyl 6-*O*-sulfation impacts glycoRNA-csRBP clustering, by generating HS2ST1 and HS6ST1 knockout U2OS cells, respectively (**Figure 4A, S4A, S4B**). Loss of 2-*O*-sulfation did not significantly alter the binding of the antibody panel (**Figure 4B**, **4C**), whereas loss of 6-*O*-sulfation reduced Siglec-11, 9D5, csDDX21, and cs-hnRNP-U bindings by 76%, 67%, 56%, and 53%, respectively (**Figure 4B**, **4C**). Sulf1 and Sulf2 are two extracellular sulfatases that can remove sulfate from the C-6 position of glucosamine of intact HS^33,34^ (**Figure 4A**). Stable expression of Sulf1 and Sulf2 (**Figure S4E**) removed total Siglec-11, 9D5, csDDX21, and cs-hnRNP-U on U2OS cells (**Figure 4B**, **4C**), suggesting that 6-*O*-sulfation of HS facilitates glycoRNA-csRBP clustering on the cell surface. Addition of exogenous HS chains with high *N*-, 6-*O*-, and 2-*O*-sulfation (rHS09) caused a loss of clustering of Siglec-11, 9D5, csDDX21, and cs-hnRNP-U, whereas the addition of HS chains with only high *N-* and 6-*O*-sulfation (rHS37) actually increased the average level of binding 2- to 3-fold) (**Figure 4B**, **4C**). The combined genetic and chemical evidence indicates that 6-*O*-sulfation of HS promotes glycoRNA-csRBP cluster formation on the cell surface.

### glycoRNA-csRBP clusters suppress VEGF-A_165_-induced signaling in primary endothelial cells

Collectively, the data suggests a biophysical mechanism underlies the HS-dependent clustering of glycoRNA-csRBP on the cell surface. To extend these studies, which were performed in tumor cell lines, to primary cells, we treated the human umbilical vein endothelial cells (HUVECs) with the pooled heparin lyases, which resulted in complete loss of binding of the antibody panel (**Figure 5A, S5A**). Treatment of HUVECs with RNases reduced binding Siglec-11, 9D5, csDDX21, and cs-hnRNP-U puncta by 73%, 83%, 96%, and 93%, respectively, without affecting 10E4 staining (**Figure 5A, S5A**). These results confirm that control of glycoRNA-csRBP clustering by HSPG is conserved between cancer cell lines and at least one primary cell model.

**Figure 5.**
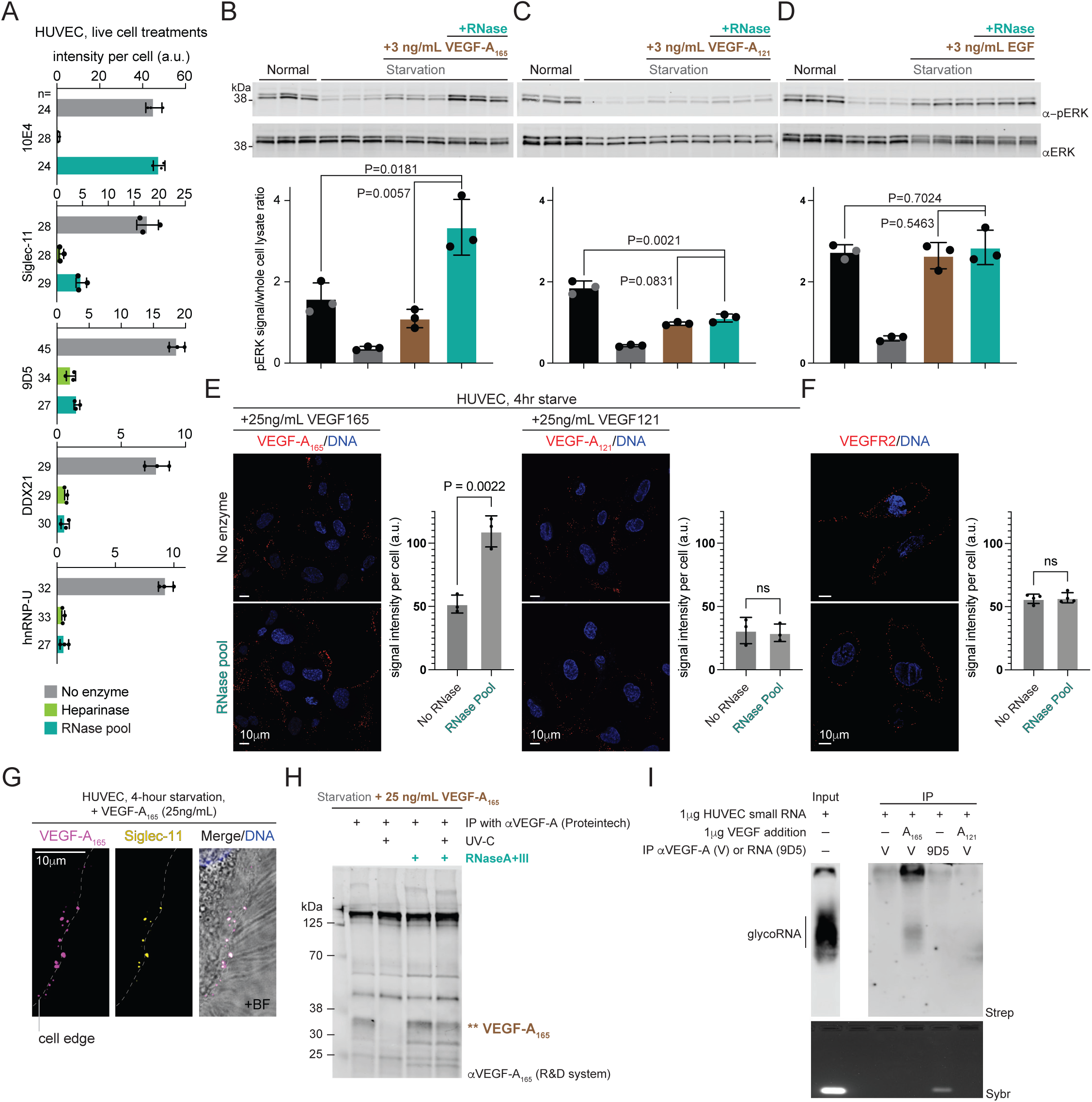
glycoRNA-csRBP clusters suppress VEGF-A_165_-induced phosphorylation of ERK in primary endothelial cells. A. Quantification of 10E4, Siglec-11, 9D5, anti-DDX21, and anti-hnRNP-U intensity per cell from 3 independent experiments. B. Western blot analysis of whole cell lysate isolated from HUVEC cells after starvation and treatment with RNase pool followed by 3 ng/mL VEGF-A_165_ stimulation. Quantification of the ratio of phosphorylated ERK (pERK) to total ERK is calculated across the biological triplicates. Statistical assessment was performed with a t-test and p values are shown. C. Western blot analysis as in (B) with HUVEC cells stimulated with 3 ng/mL of VEGF-A_121_. D. Western blot analysis as in (B) with HUVEC cells stimulated with 3 ng/mL of EGF. E. Representative confocal images of HUVEC cells after serum starvation, treatment with or without RNase pool, and the treatment with VEGF-A_165_ or VEGF-A_121_, finally imaging with anti-VEGF-A_165_ (red) or anti-VEGF-A (red). DNA was stained with DAPI (blue). Scale bar, 10 µm. Quantification of the images with number of cells noted per biological triplicate. Statistical assessment was performed with a t-test and p values are shown. F. Representative confocal images of HUVEC cells with the treatment with or without RNase pool, and imaging with anti-VEGFR2 (red). DNA was stained with DAPI (blue). Scale bar, 10 µm. Quantification of the images with number of cells noted per biological triplicate. Statistical assessment was performed with a t-test and p values are shown. G. Representative confocal images of HUVEC cells after serum starvation and treated with or without VEGF-A_165_ and then costained with anti-VEGF-A_165_ (purple) and Siglec-11 (yellow). DNA was stained with DAPI (blue). Zoomed region shown as an inset. Scale bar, 10 µm. H. Lysates from HUVEC cells after serum starvation, treatment with 25 ng/mL VEGF-A_165_, and UV-crosslinking, were treated with or without RNase pool and immunoprecipitated (IP) with an anti-VEGFA antibody (Proteintech). Immunoprecipitated samples were analyzed by Western blot using an anti-VEGF-A_165_ antibody (R&D system). I. 1 µg of VEGF-A_165_ or VEGF-A_121_ was immunoprecipitated with an anti-VEGFA antibody. 1 µg of HUVEC small RNA was then incubated to beads preconjugated to anti-VEGF-A or 9D5 antibodies. RNA was extracted from the beads, purified, rPAL labeled, and finally analyzed by Northern blot. 1 µg of HUVEC small RNA served as input of glycoRNA.

The colocalization of HS with glycoRNA-csRBPs suggested the possibility that growth factor signaling normally thought to depend on HS action as a coreceptor might in fact be modulated by glycoRNA-csRBP clusters. To test this hypothesis, we serum starvation HUVECs and used western blotting to detect total and phosphorylated ERK (ERK and pERK) after VEGF-A stimulation. Various proteoforms of VEGF-A including VEGF-A_121_ and VEGF-A_165_ bind the VEGFR via their N-terminal domains; the extended C-terminus of VEGF-A_165_ enables interactions with heparan sulfates and neuropilin-1 (a VEGFR coreceptor)^35,36^. Serum starvation of the HUVECs reduced phosphorylation of ERK relative to total ERK by ∼77% (**Figure 5B**); upon stimulation with 3 ng/mL of VEGF-A_165_ or VEGF-A_121_, we observed >50% recovery of the pERK levels (**Figure 5B, brown bars**). Pre-treatment of the cells with the RNase pool before the addition of VEGF-A_165_ resulted in a 3-fold and 2.1-fold increase in pERK compared to cells without RNase treatment and to homeostatic levels in untreated cells, respectively (**Figure 5C**). This was not true for VEGF-A_121_ which showed no change in the recovery pERK levels with or without RNase pre-treatment (**Figure 5C**). Outside of VEGF-A, other growth factors like epidermal growth factor (EGF) can also signal through the ERK pathway; however EGF is not a HSPG-binding protein^37^. EGF (3 ng/mL) was able to fully restore pERK to pre-starved levels and was similarly insensitive to RNase as VEGF-A_121_ (**Figure 5D**). The RNase-dependent effect of VEGF-A_165_ (and the insensitivity of VEGF-A_121_ and EGF) are also seen at 25 ng/mL of each growth factor demonstrating robustness of this mechanism across a range of concentrations (**Figure S5B-S5D**).

We next assessed how the loss of cell surface RNA directly impacts cell surface association of VEGF-A. Serum starvation and pre-treatment with the RNase pool led to ∼2x more binding of VEGF-A_165_ to the cell surface while VEGF-A_121_ saw no changes on its cell surface association (**Figure 5E**). To determine if this was related to change only in VEGF-A_165_ or its receptor on HUVECs VEGFR2^38,39^, we imaged VEGFR2 under the same conditions and found no changes in the receptor abundance on the cell surface (**Figure 5F**). To further explore the mechanism of enhanced signal transduction after the loss of cell surface RNA, we examined the spatial relationship between VEGF-A_165_ and glycoRNA-csRBP clusters. Co-staining of VEGF-A_165_ and Siglec-11 on HUVECs revealed 78% of Siglec-11 puncta overlapped with VEGF-A_165_ and 40% of VEGF-A_165_ puncta overlapped with Siglec-11 (**Figure 5G, S5E, S5F**). VEGF-A_165_ was absent on cells without the exogenous addition of VEGF-A, while Siglec-11 clusters were clearly present (**Figure S5E**).

Finally, to understand how the cell surface RNA itself could repress the ability for VEGF-A_165_ to bind the cell surface, we investigated if VEGF-A_165_ directly interacts with cell surface RNA. After starvation and VEGF-A_165_ addition, cells were UV-C crosslinked to produce covalent bonds between directly bound RNA-protein complex and immunoprecipitates of the bound VEGF-A_165_ were evaluated. Without UV-C we were able to recover VEGF-A_165_ however upon crosslinking the native migrating band disappears (**Figure 5H**). We predicted this was due to covalent UV-crosslinking of VEGF-A_165_ to RNA and consistent with this, RNase treating the lysate after UV-C exposure restores the native migrating VEGF-A_165_ band (**Figure 5H**). Next, we tested if VEGF-A_165_ could directly interact with glycoRNAs. Co-incubating VEGF-A_165_ with small RNA from HUVEC cells and subsequent VEGF-A IP and rPAL labeling resulted in selective capture of glycoRNAs (**Figure 5I**). If we omitted VEGF-A_165_ or added VEGF-A_121_ we were unable to isolate glycoRNAs (**Figure 5I**). Further, the anti-RNA antibody 9D5 was able to capture bulk small RNA (Sybr signal) but unable to preferentially isolate glycoRNA species in vitro like we observe when using VEGF-A_165_ to capture small RNA (**Figure 5I**), indicating that VEGF-A_165_ selectively interacts with glycoRNAs.

## Discussion

Heparan sulfate proteoglycans (HSPGs) are widely appreciated to govern critical processes at the cell surface in health and disease. Here, we begin to refine this view with the characterization of how glycoRNAs and RBPs assemble into clusters nucleated around HSPGs. Through genetic and enzymatic perturbation, we demonstrate that the extended HS chains are important for the clustering and if removed, repopulate the cell surface first, suggesting that the glycoRNA-csRBPs assemble on or around already presented HSPGs. Molecularly, the sulfation of the HS chains is critical, with *N*- and 6-*O*-sulfation responsible for promoting the glycoRNA-csRBP clustering. Finally, we establish a new mechanistic pathway for information to be transferred from outside to inside of a cell: through the direct binding and regulation of growth factors like VEGF-A_165_ to cell surface glycoRNA.

Our genome-wide knockout screening efforts provided the initial evidence for HS-regulation of glycoRNA-csRBP clusters; however it also highlights the critical role that RNAs on the cell surface play in the landscape of endogenous ligands for Siglec-11. Comparing the highly enriched genes that caused reduced Siglec-11 binding there was much stronger overlap to a general anti-RNA antibody than a well characterized sialic acid-binding lectin (MAA-I). This motivates a more detailed examination of the bona fide ligands of Siglec-11, their biophysical association with Siglec-11, and the regulatory role they could in the interaction of cells that express Siglec-11 such as macrophages or microglia.

Considering the HSPGs as the nucleating site for glycoRNA-csRBPs, the clustering mechanism remains to be established. Perhaps the RBPs can act as a bridge that bind to glycoRNAs and to HS chains of HSPGs. It also remains unclear whether these clusters depend on a specific HSPG, given that cells typically express multiple cell surface proteoglycans. The observation that several genes involved in glycosylphosphatidylinositol biosynthesis were discovered in the genome wide CRISPR screen suggests that one or more GPI-anchored proteoglycans (glypicans) might be part of the glycoRNA-csRBP complexes.

Finally, RNA’s role in regulating biological processes is expansive and spans its catalytic, scaffolding, and information-carrying capabilities^40–42^, however these functions have traditionally been restricted to intracellular pathways. Having evidence that VEGF-A_165_ binds at glycoRNA-csRBP clusters and that its binding is sensitive to RNase suggests a physical role of RNA in the organization of cell surface domains. Further and perhaps most interestingly, our data demonstrate that canonical trans-membrane signal transduction by VEGF-A_165_ is modulated by the presence of cell surface RNA. Critically, our in vitro data demonstrate selective regulation of signaling by glycoRNAs by VEGF-A_165_ but not VEGF-A_121_, suggesting that the expression of specific isoforms of VEGF-A could directly target cell surface glycoRNAs. Together, our data establishes a new type of RNA regulation where the abundance or organization of topologically extracellular RNAs can directly modulate intracellular signaling cascades, impacting cellular decision making.

---

**Figure S1.**
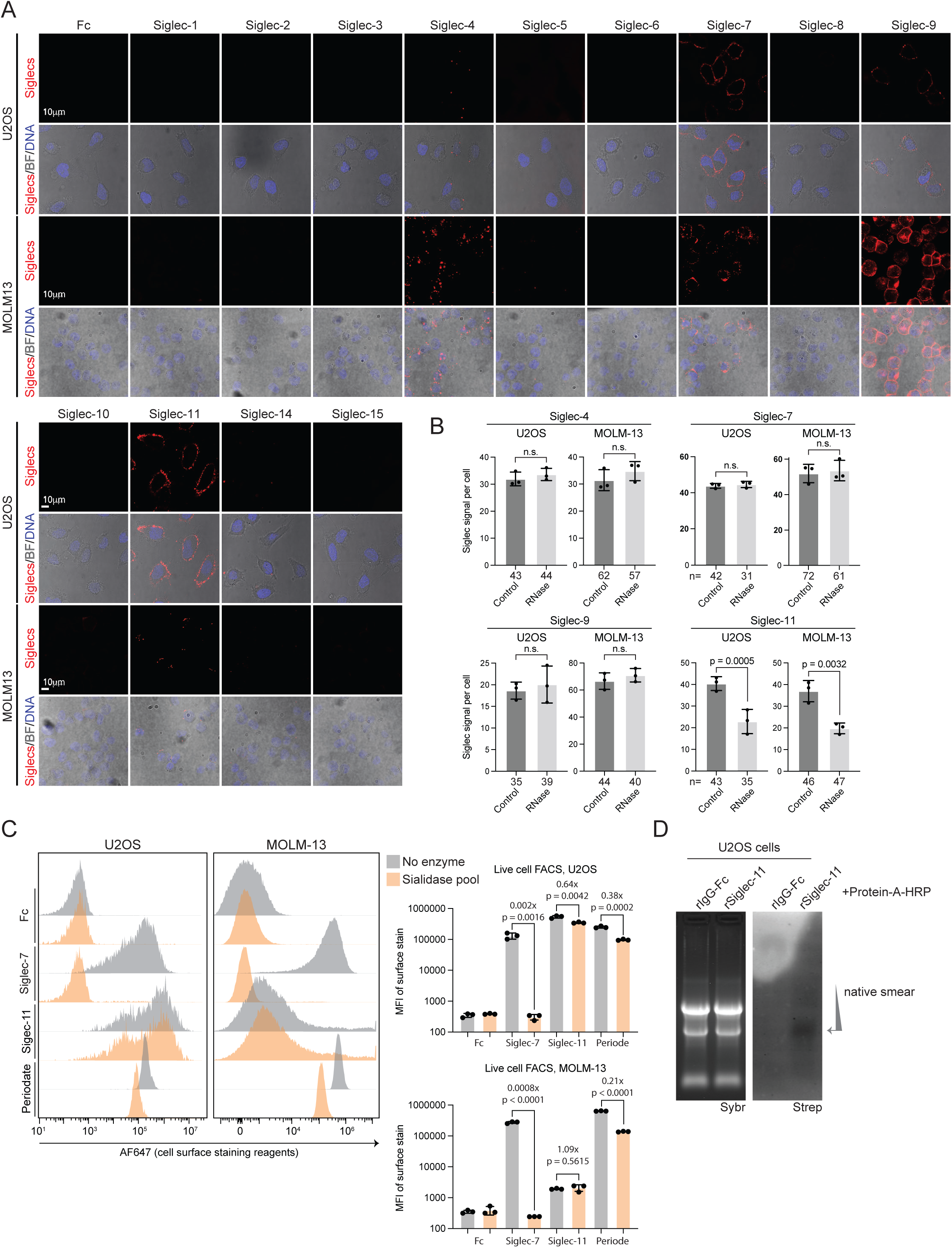
Binding profile and RNase-dependency of 13 human Siglec receptors. A. Representative confocal images of U2OS and MOLM-13 cells stained live with IgG1 or the indicated Siglec-Fc reagents (red). DNA was stained with DAPI (blue). Scale bar, 10 µm. B. Quantification of the indicated Siglecs intensity in U2OS and MOLM-13 cells treated with RNase pool for 30 min from 3 independent experiments with the number (n) of cells analyzed noted. C. Representative histograms (left) from flow cytometry experiments of U2OS or MOLM-13 cells treated live with or without a sialidase pool and then analyzed for surface binding of Fc-IgG (control), Siglec-7, Siglec-11, or total surface glycan (periodate) signal. Triplicate experiments are quantified with the mean fluorescence intensity (MFI) on the right for each cell line, and t-tests were used to assess statistical differences, with the fold changes noted. D. RNA blotting as in Figure 1E, here on U2OS cells.

**Figure S2.**
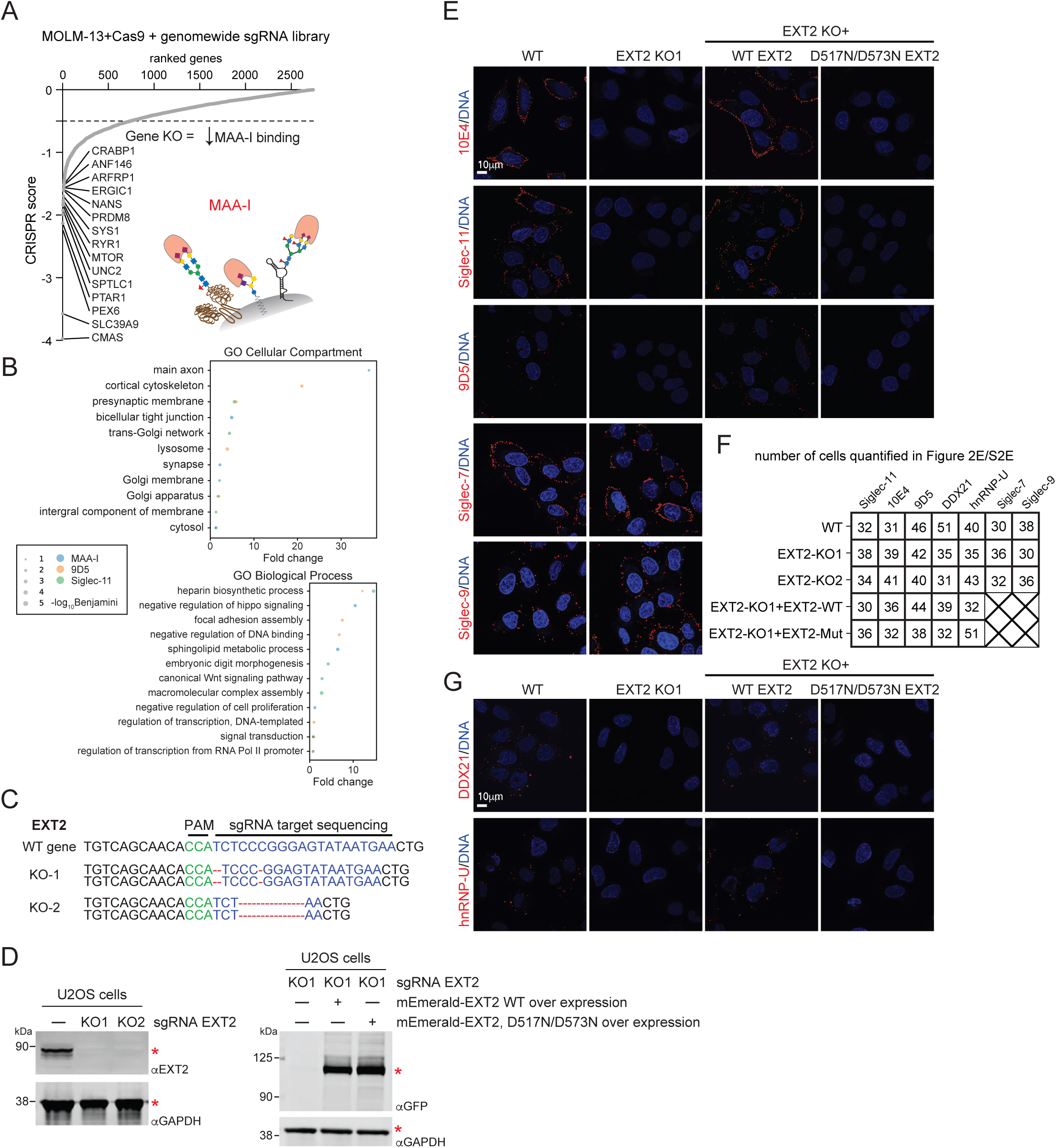
EXT2 knockouts and rescues modulate glycoRNA-csRBP clusters. A. Dot plot of genes identified in the genome-wide CRISPR knockout (KO) screen for loss of MAA-I cell surface binding ranked by CRISPR score. The top 15 gene names are displayed with a line drawn at the -0.8 score cut off. The inset cartoon illustrates how MAA-I could interact with a cell surface glycoRNA. B. Gene ontology (GO) cellular compartment (top) and biological process (bottom) analysis of KO screen hits from MAA-I (blue), 9D5 (red), and Siglec-11 (green). The top 4 terms across the three screens were intersected and the union is displayed with the significance of each term represented by circle size and plotted on the x-axis by their fold enrichment. C. Sequence alignment of partial EXT2 coding sequences from WT and EXT2 KO U2OS cells. Protospacer adjacent motif (PAM), green; sgRNA target, blue; mutated sequence, red. D. Western blot analysis of whole cell lysate. (left) from wild-type (WT) and EXT2 knock-out U2OS cell lines and (right) EXT2-KO1, KO-mEmerald-EXT2, EXT2 KO-mEmerald-D517N/D573N EXT2. In both, GAPDH served as loading control. E. Representative confocal images of WT, and EXT2 KO U2OS cells stained with Siglec-7 and Siglec-9 (red), separately. DNA was stained with DAPI (blue). Scale bar, 10 µm. F. Number of cells quantified in Figure 2D, 2F, S2E, S2G across 3 independent experiments. G. Representative confocal images of EXT2 KO-mEmerald-EXT2 and EXT2 KO-mEmerald-D517N/D573N EXT2 U2OS cells. DNA was stained with DAPI (blue). Scale bar, 10 µm.

**Figure S3.**
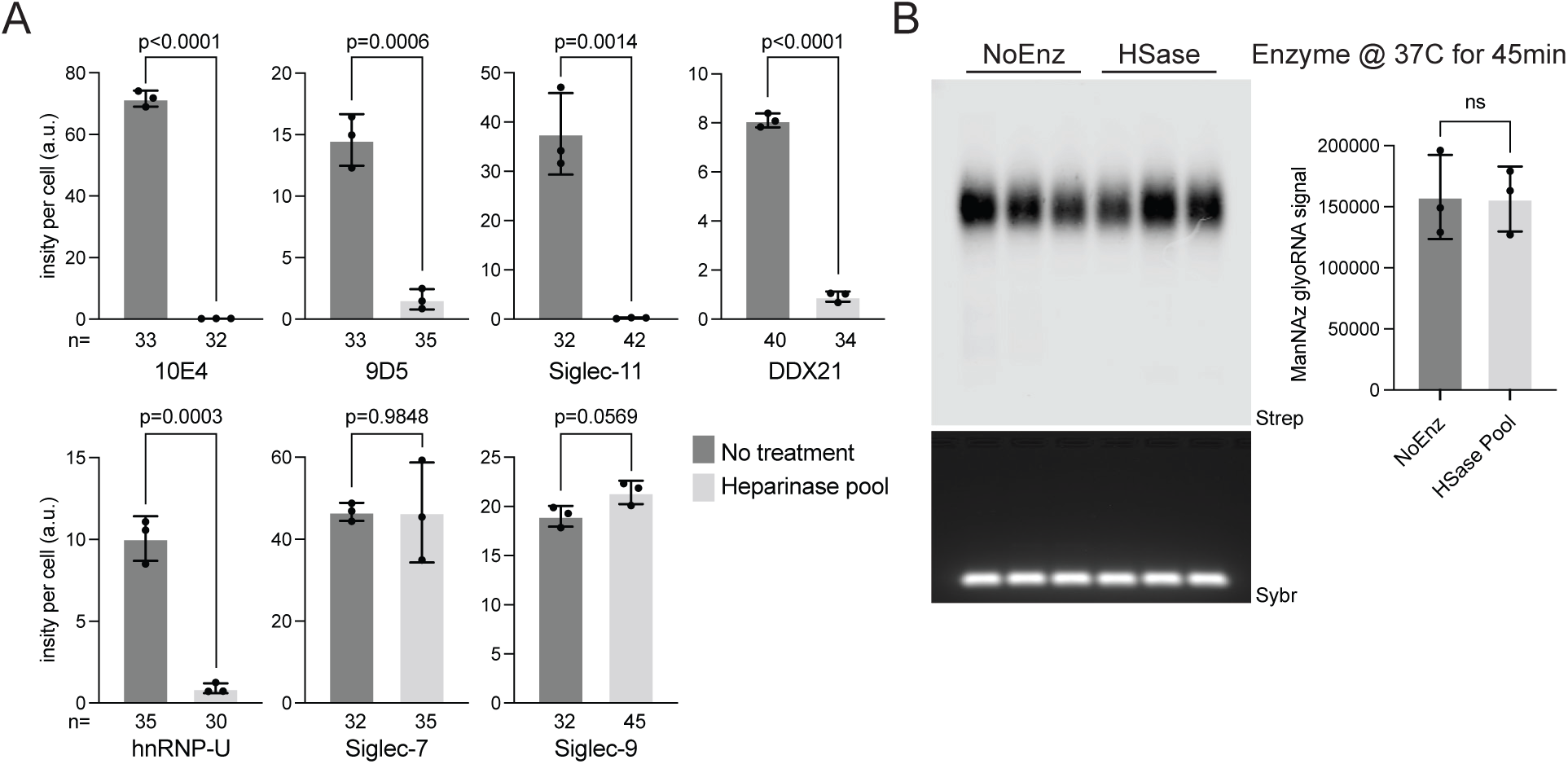
Quantification of cell surface ligands after heparinase treatment. A. Quantification of data in Figure 3A with the number of cells per condition from 3 independent experiments is shown. Statistical assessment was performed with a t-test and p values are shown. B. RNA blotting of U2OS small RNA labeled with Ac_4_ManNAz detected DBCO-biotin and in vitro digested with no enzyme or heparinase pool. small RNA (Sybr, bottom) and biotin detection (Strep, top) is shown. Quantification plotted on the right and statistical assessment of the intensities was performed with a t-test.

**Figure S4.**
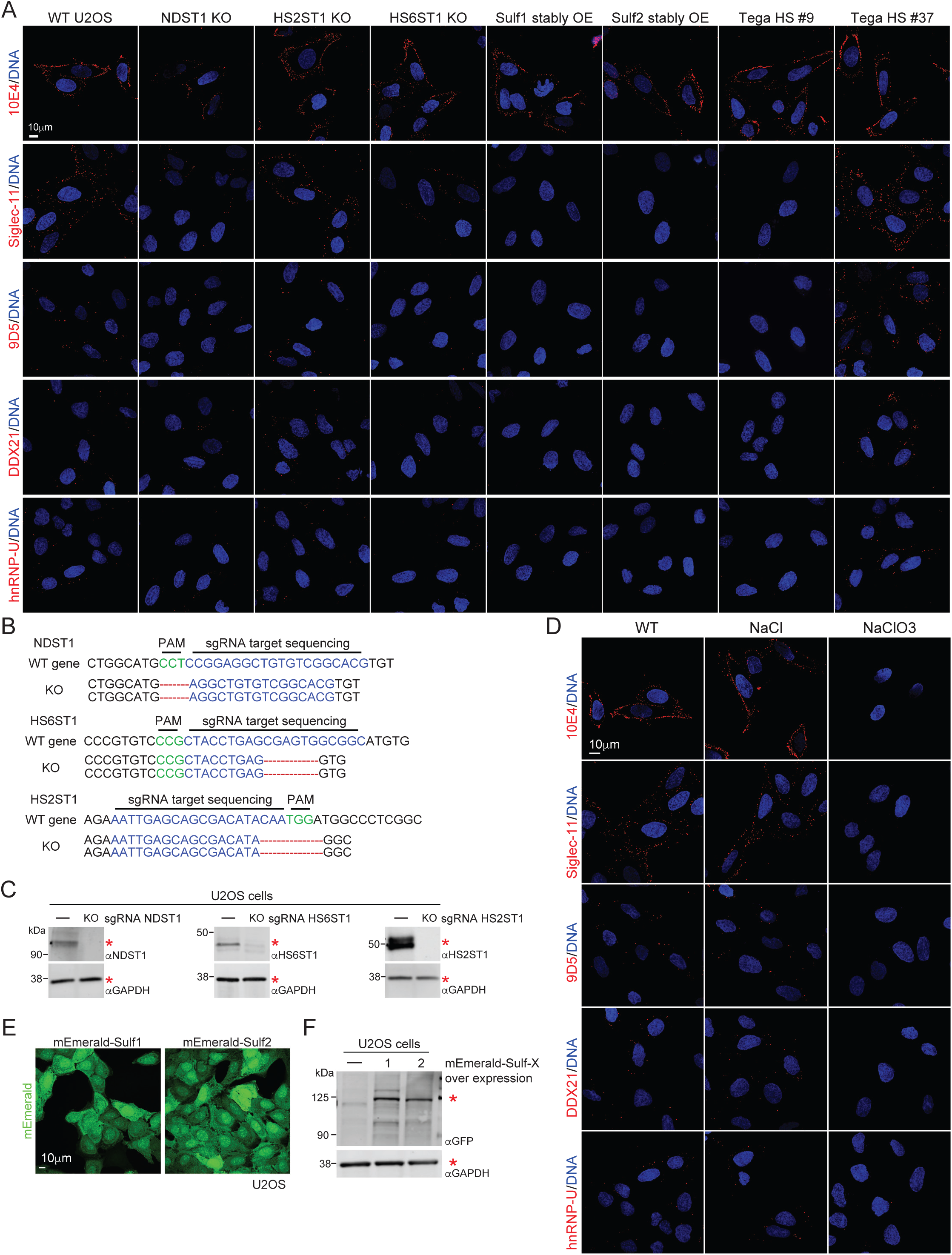
Effects of cellular sulfation and sulfatases on glycoRNA-csRBP clustering. A. Sequence alignment of partial NDST1, HS6ST1, and HS2ST1 coding sequences from wild-type (WT) and the indicated knock-out (KO) U2OS cell lines. Protospacer adjacent motif (PAM), green; sgRNA target, blue; mutated sequence, red. B. Western blot analysis of whole cell lysate from WT, NDST1 KO, HS6ST1, and HS2ST1 KO U2OS cells. GAPDH served as loading control. C. Representative confocal images of WT and NaCl or NaClO_3_ treated U2OS cells stained with 10E4, Siglec-11, 9D5, anti-DDX21, and anti-hnRNP-U (all in red). DNA was stained with DAPI (blue). Scale bar, 10 µm. D. Representative confocal images of mEmerald-Sufl1 and mEmerald-Sulf2 stably overexpressing U2OS cells. DNA was stained with DAPI (blue). Scale bar, 10 µm. E. Western blot analysis of whole cell lysate isolated from mEmerald-Sufl1 and mEmerald-Sulf2 stably overexpressing U2OS cells. GAPDH served as loading control.

**Figure S5.**
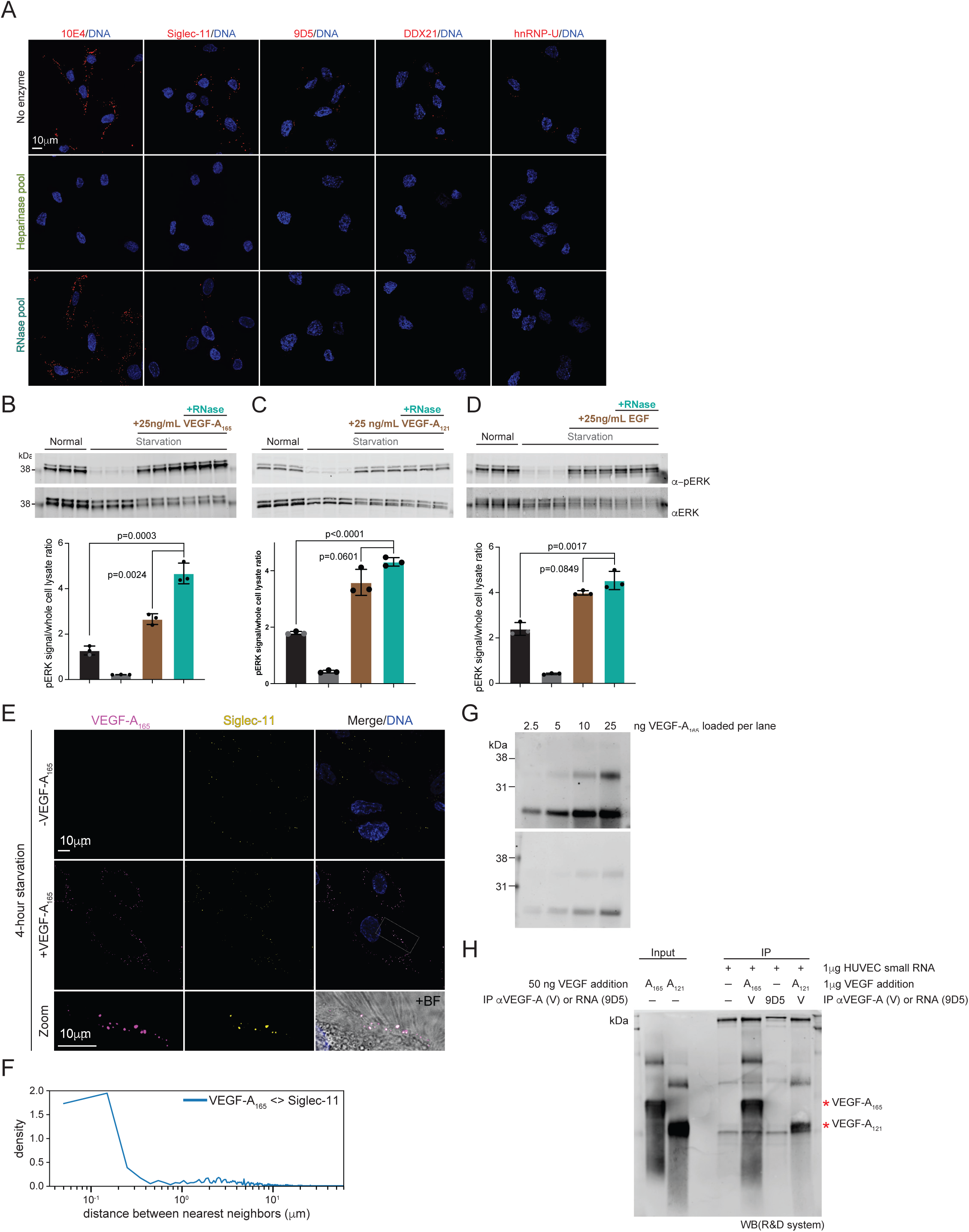
HUVEC response to live cell enzymes, VEGF-A, and EGF addition. A. Representative confocal images of HUVEC cells treated with the RNase pool or heparinase pool, and stained live with 10E4, Siglec-11, 9D5, anti-DDX21, and anti-hnRNP-U (all in red). DNA was stained with DAPI (blue). Scale bar, 10 µm. B. Western blot analysis of whole cell lysate isolated from HUVEC cells after starvation and treatment with RNase pool followed by 25 ng/mL VEGF-A_165_ stimulation. Quantification of the ratio of phosphorylated ERK (pERK) to total ERK is calculated across the biological triplicates. Statistical assessment was performed with a t-test and p values are shown. C. Western blot analysis as in (B) with HUVEC cells stimulated with 25 ng/mL of VEGF-A_121_. D. Western blot analysis as in (B) with HUVEC cells stimulated with 25 ng/mL of EGF. E. Representative confocal images of HUVEC cells after serum starvation and treated with or without VEGF-A_165_ and then costained with anti-VEGF-A_165_ (purple) and Siglec-11 (yellow). DNA was stained with DAPI (blue). Zoomed region shown as an inset. Scale bar, 10 µm. F. Nearest neighbor distance analysis of the VEGF-A_165_ and Siglec-11 in Figure S5E. For each pair, the nm distance from VEGF-A_165_ to Siglec-11 was calculated across. These values were plotted in a density histogram. G. Western blot analysis of the indicated amount of VEGF-A_165_ or VEGF-A_121_. H. 1 µg of VEGF-A_165_ or VEGF-A_121_ was immunoprecipitated with an anti-VEGFA antibody (Proteintech). Immunoprecipitated samples were analyzed by Western blot using an anti-VEGF-A_165_ antibody (R&D system).

## Acknowledgments

We thank Kayvon Pedram, Carolyn R. Bertozzi, Eliezer Calo, Reese Caldwell, and other members of the Flynn Lab for helpful comments and discussions. This work was supported by grants from Wellcome Trust (K.T. RG83195, G106133), the UKRI Medical Research Council (K.T. RG83195) and Leukaemia UK (K.T. G117699), Burroughs Wellcome Fund Career Award for Medical Scientists (R.A.F.), the Sontag Foundation Distinguished Scientist Award (R.A.F.), the Rita Allen Foundation (R.A.F.), and the Scleroderma Research Foundation (R.A.F.).

## Author Contributions

R.A.F. conceived and supervised the project. P.C. designed and performed the bulk of the experiments. L.K. and C.G.L. performed the flow cytometry experiments and analysis. J.P. designed the csRBP experiments. K.T., E.Y., J.M.L.D., and R.A.F. designed, performed and analyzed the knockout screens. J.E. designed inhibitor and knockout experiments surrounding heparan sulfate pathways. R.A.F. and P.C. wrote the manuscript. All authors discussed the results and revised the manuscript.

## Methods

### Cell culture

MOLM-13 cells were cultured in Roswell Park Memorial Institute 1640 Medium (Thermo Fisher Scientific). U2OS (ATCC) were cultured in McCoy’s 5A Medium (Thermo Fisher Scientific). MOLM-13 and U2OS cells were supplemented with 10% fetal bovine serum (FBS, Thermo Fisher Scientific) and 1x Penicillin/Streptomycin (Thermo Fisher Scientific). Primary umbilical vein endothelial cells (HUVEC) (ATCC) were cultured in Vascular Cell Basal Medium (ATCC) supplemented with Endothelial Cell Growth Kit-VEGF (ATCC). All the experiments on HUVEC are performed before passage 10. All cells were cultured at 37°C with 5% CO_2_ and maintained as mycoplasma negative. U2OS cells were transfected with Avalanche-Omni Transfection Reagent (EZ Biosystems).

### Live cell enzyme and chemical treatments

For RNase treatment, RNase A (Sigma) and ShortCut RNase III (New England Biolabs, NEB) were added directly to the cell culture at a final concentration of 18 µM and 100 U/mL, separately. MOLM-13, U2OS, and HUVEC cells were all treated for 45 minutes.

For Ac_4_ManNAz labeling, stocks of N-azidoacetylmannosamine-tetraacylated (Ac_4_ManNAz, Click Chemistry Tools) were made to 500 mM in sterile dimethyl sulfoxide (DMSO). Ac4ManNAz was added to the cell culture at a final concentration of 100 μM for 24 hours before collection.

For heparinase treatment, heparinase I (NEB), heparinase II (NEB), and heparinase III (NEB) were added directly to the cell culture at a final concentration of 4 units/mL (each) for 30 minutes.

For sodium chloride (NaCl) and sodium chlorate (NaClO_3_) treatment, NaCl and NaClO3 were added to the cell culture at a final concentration of 50 mM for 24 hours.

For exogenous Tega heparan sulfate chain treatment, rHS09 (TEGA Therapeutics) and rHS37 (TEGA Therapeutics) were added directly to the cell culture at a final concentration of 4 µM for 60 minutes.

For serum starvation, HUVECs were cultured in Vascular Cell Basal Medium (Bioresource Center, PCS100030) without Endothelial Cell Growth Kit-VEGF (Biosource Center) after being washed briefly for 3 times with phosphate buffer saline (PBS). VEGF-A_165_ (Thermo Fisher Scientific), VEGF-A_121_ (Thermo Fisher Scientific) and EGF (Thermo Fisher Scientific) were then added to starved HUVECs at a final concentration of 3 ng/mL and 25 ng/mL for 5 minutes, separately.

### Live cell labeling, confocal microscopy, quantification and statistical analysis

Adherent cells were cultured on glass coverslips # 1.5 (Bioscience Tools) 24 hours before labeling. MOLM-13 cells were counted and then blocked as per the manufacturer’s protocol with Human TruStain FcX (Fc block, BioLegend) for 15 minutes on ice before labeling. For Siglecs staining in live cells, 1 µg/mL of recombinant human IgG1 Fc (R&D Systems), Siglec-1 Fc chimera protein (R&D Systems), Siglec-2 Fc chimera protein (R&D Systems), Siglec-3 Fc chimera protein (R&D Systems), Siglec-4 Fc chimera protein (R&D Systems), Siglec-5 Fc chimera protein (R&D Systems), Siglec-6 Fc chimera protein (R&D Systems), Siglec-7 Fc chimera protein (R&D Systems), Siglec-8 Fc chimera protein (R&D Systems), Siglec-9 Fc chimera protein (R&D Systems), Siglec-10 Fc chimera protein (R&D Systems), Siglec-11 Fc chimera protein (R&D Systems), Siglec-14 Fc chimera protein (R&D Systems), and Siglec-15 Fc chimera protein (R&D Systems) were precomplexed with 0.5 µg/mL of donkey anti-human IgG AF647 (ImmunoResearch) secondary antibody in FACS buffer (0.5% BSA (Sigma) in 1x PBS) for 45 minutes on ice. For 9D5 and 10E4 staining, 2.5 µg/mL of 9D5 (Absolute Antibody), 1 µg/mL of anti-heparan sulfate 10E4 (amsbio), and anti-VEGF (R&D Systems) were precomplexed with 1.25 µg/mL of goat anti-Rabbit AF647 secondary antibody (ThermoFisher Scientific), 0.5 µg/mL of goat anti-Mouse AF647 secondary antibody (ThermoFisher Scientific), or donkey anti-Goat IgG AF647 (ThermoFisher Scientific) secondary antibodies in FACS buffer for 45 minutes on ice, separately. For VEGF-A_165_, VEGF-A_121_, and VEGFR2 staining, 1 µg/mL of anti-VEGF-A_165_ (R&D Systems), 1 µg/mL of anti-VEGFA (Proteintech), and 1 µg/mL anti-VEGFR2 (R&D Systems) were precomplexed with 0.5 µg/mL of goat anti-Rabbit AF647 secondary antibody and donkey anti-Goat AF647 secondary antibody (ThermoFisher Scientific). Precomplexed antibodies were then incubated with cells for 45 minutes on ice. For DDX21 and hnRNP-U staining, 2.5 µg/mL of anti-DDX21 (Novus Biological) and anti-hnRNP-U (Proteintech) were incubated with cells for 45 minutes on ice. Cells were gently washed twice by FACS buffer and then stained with 2.5 µg/mL of goat anti-Rabbit AF647 for 30 minutes on ice.

For 9D5 and Siglecs co-staining, recombinant human Siglec Fc chimera proteins were precomplexed with donkey anti-Human IgG AF488 (ImmunoResearch), 9D5 was precomplexed with goat anti-Rabbit AF647 secondary antibody. For 9D5, Siglec-11, and 10E4 co-staining, 10E4 was precomplexed with goat anti-mouse AF568 (ThermoFisher Scientific). For VEGF-A_165_ and Siglec-11 co-staining, anti-VEGF-A_165_ was precomplexed with donkey anti-Goat AF647 secondary antibody, recombinant human Siglec-11 Fc chimera protein was precomplexed with donkey anti-Human IgG AF488. Antibodies or regents were precomplexed in separate tubes with the same concentration as used in single channel staining and then mixed before adding to cells. After staining, cells were washed three times with ice-cold PBS and a fixation was performed with 3.7% formaldehyde for 15 minutes at room temperature in the dark. Nuclei were stained with 0.1 µg/mL DAPI in PBS. MOLM-13 cells were applied to glass slides using a CytoSpin centrifuge (ThermoFisher Scientific) at 500x g for 5 minutes.

For mEmerald-positive U2OS cell imaging, cells were fixed with 3.7% formaldehyde for 15 minutes at room temperature and then stained with DAPI. Finally, all samples above were mounted in ProLong Diamond Antifade Mountant (ThermoFisher Scientific) and a coverglass was sealed over the samples with nail polish. All samples were then imaged on a Leica SP8 STED ONE microscope with 63x oil lens. Images were acquired using Leica LAS X software. The DAPI channel was acquired with a PMT detector while all other channels were imaged using Hybrid detectors.

For quantification and statistical analysis, at least three random regions of interest (ROIs) from three independent samples were acquired across one or more z-slices. To analyze the colocalization, images were processed using Imaris Microscopy Image Analysis software (Oxford Instruments). A single z-slice from each ROI was taken, selected to be near the middle of the cells (with respect to their z-thickness), and the spot-finder function was used to identify spots of roughly 0.5 µm. In-software background subtraction was used as the default settings, and spots were selected by thresholding spot quality at the elbow of the distribution. This resulted in a series of x- and y-positions for each spot from each channel, which were then exported for quantitative analysis. Colocalization of spots from paired channels were analyzed by implementing a custom Python script (https://github.com/FlynnLab/jonperr) to identify the nearest neighbors of each spot (in nanometers, nm) with a k-d tree algorithm (scipy.spatial.KDTree). Then, the distances between nearest neighbors were calculated for each pair of targets across all ROIs and plotted in a histogram. To assess the relative fraction of each spot type (channel #1) within the other pair’s spots (channel #2), we calculated a Manders’ colocalization coefficient (MCC) using the aforementioned Python script. We performed this calculation in both directions: spots of channel #1 in total channel #2 spots, and the reverse.

To quantify and compare the intensities of spots on the cell surface, Leica LAS X software was used to identify ROIs throughout the entire 4x slice z-stack. To quantify and compare the spot numbers of spots on the cell surface, Imaris was used to identify spots throughout the entire 4x slice z-stack. The mean intensities and numbers of ROIs were then divided by the cell numbers and compared across groups. Statistical analysis and data plotting were performed using GraphPad Prism 10.

### Live cell flow cytometry

MOLM-13 cells were directly counted. U2OS cells were gently lifted with Accutase (Sigma-Aldrich) for 3 minutes at 37°C, quenched with growth media, and then counted. For each condition, 50,000 cells were used. For antibody staining, MOLM-13 cells were blocked as per the manufacturer’s protocol with Human TruStain FcX in FACS buffer for 15 minutes on ice. 1 µg/mL of recombinant human IgG1 Fc, Siglec-7 Fc chimera protein, and Siglec-11 Fc chimera protein were precomplexed with 0.5 µg/mL of donkey anti-human IgG AF647 secondary antibody in FACS buffer for 45 minutes on ice. Precomplexed antibodies were then added to bind cells on ice for 45 minutes. For live cell periodate labeling of cell surface glycans, cells were washed twice with cold PBS + Ca + Mg and then incubated at 4°C in cold PBS + 1 mM sodium periodate for 5 minutes at 1 million cells per mL. Cells were then quenched with 1 mM glycerol added to the PBS, and then cells were washed twice with cold PBS. Cells were then incubated at 4°C in cold FACS buffer + 25 µM aminooxy-biotin (Cayman Chemical) + 10 mM aniline for 30 minutes at 1 million cells per mL. Cells were mixed via pipetting halfway through the incubation. Cells were then washed once with cold 1x PBS and blocked as per the manufacturer’s protocol with Human TruStain FcX in FACS buffer for 15 minutes on ice. Cells were then stained for 30 minutes on ice with Strep-AF647 at 1 µg/mL. After staining, all cells were spun at 4°C for 3 minutes at 400x g and supernatant discarded. Cells were washed once with 150 µL of FACS buffer, spun under the same conditions, and finally resuspended in FACS buffer containing 0.1 µg/mL DAPI. Data collection occurred on a BD Biosciences LSRFortessa 3 and a gating strategy was used to isolate live, single cell, to examine antibody binding using FlowJo Software (FlowJo LLC).

### Live cell RNA proximity labeling

Samples were prepared similarly to the flow cytometry workflow as described above however rather than dye-conjugated secondaries, here horseradish peroxidase (HRP) conjugates secondaries were used. 1 µg/mL of recombinant human IgG1 Fc or Siglec-11 Fc chimera protein was precomplexed with 0.5 µg/mL of Protein A -HRP (Cell Signaling Technology) on ice for 30 minutes. Cells were adjusted to 1 million cells per mL of FACS and then the precomplexed antibodies were added for staining. Staining occurred for 60 minutes at 4°C on rotation, after which cells were pelleted, supernatants discarded and cells washed once in ice-cold PBS. This wash is important to remove excess BSA in the FACS buffer. Next, cells were gently but quickly resuspended in 985 µL of 200 µM biotin-aniline (Iris Biotech) in PBS at 25°C. To this, 15 µL of 100 mM H2O2 was quickly added, tubes capped and inverted, and the reaction allowed to proceed for 2 minutes at 25°C. Precisely after 2 minutes, the samples were quenched by adding FACS buffer with sodium azide and sodium ascorbate to a final concentration of 5 mM and 10 mM, respectively. Samples were inverted and pelleted at 4°C. Cell pellets were directly lysed in 500 µL of RNAzol RT (Molecular Research Center). Samples were shaken at 50°C for 5 minutes. To phase separate the RNA, 0.4X volumes of water was added, vortexed, let to stand for 5 minutes at 25°C and lastly spun at 12,000x g at 4°C for 15 minutes. The aqueous phase was transferred to clean tubes and 1.1X volumes of isopropanol was added. The RNA was then purified over a Zymo column (Zymo Research). For all column cleanups, we followed the following protocol. First, 350 µL of pure water was added to each column and spun at 10,000x g for 30 seconds, and the flowthrough was discarded. Next, precipitated RNA from the RNAzol RT extraction (or binding buffer precipitated RNA, below) was added to the columns, spun at 10,000x g for 20 seconds, and the flowthrough was discarded. This step was repeated until all the precipitated RNA was passed over the column once. Next, the column was washed three times total: once using 400 µL of RNA Prep Buffer (3M GuHCl in 80% EtOH), twice with 400 µL of 80% ethanol. The first two spins were at 10,000x g for 20 seconds, the last for 30 seconds. The RNA was then treated with Proteinase K (Ambion) on the column. Proteinase K is diluted 1:19 in water and added directly to the column matrix and then allowed to incubate on the column at 37°C for 45 minutes. The column top was sealed with either a cap or parafilm to avoid evaporation. After the digestion, the columns were brought to room temperature for 5 minutes; lowering the temperature is important before proceeding. Next, eluted RNA was spun out into fresh tubes and a second elution with water was performed. To the eluate, 1.5 µg of the mucinase StcE (Sigma-Aldrich) was added for every 50 µL of RNA, and placed at 37°C for 30 minutes to digest. The RNA was then cleaned up again using a Zymo column. Here, 2X RNA Binding buffer (Zymo Research) was added and vortexed for 10 seconds, and then 2X (samples + buffer) of 100% ethanol was added and vortexed for 10 seconds. The final RNA was quantified using a Nanodrop. In vitro RNase or Sialidase digestions took place by digesting 50 µg total RNA with either, nothing, 4 µL RNase Cocktail (ThermoFisher Scientific), or 4 µL of α2-3,6,8,9 Neuraminidase A (NEB,) in 1x NEB Glyco Buffer #1 (NEB) for 60 minutes at 37°C. After digestion, RNA was purified using a Zymo column as noted above and was then ready for gel analysis.

In order to visualize the labeled RNA, the samples were run on a denaturing agarose gel, transferred to nitrocellulose membranes (Bio-Rad Laboratories), stained with IRDye 800CW Streptavidin (LI-COR Biosciences). After elution from the column as described above, the RNA is combined with 12 µL of Gel Loading Buffer II (GLBII, 95% formamide, 18 mM EDTA, 0.025% SDS) with a final concentration of 1x SybrGold (ThermoFisher Scientific) and denatured at 55°C for 10 minutes. It is important to not use GLBII with dyes. Immediately after this incubation, the RNA is placed on ice for at least 2 minutes. The samples were then loaded into a 1% agarose, 0.75% formaldehyde, 1.5x MOPS buffer (Lonza) denaturing gel. Precise and consistent pouring of these gels is critical to ensure a similar thickness of the gel for accurate transfer conditions; we aim for approximately 1 cm thick of solidified gel. RNA was electrophoresed in 1x MOPS at 115V for between 34 or 45 minutes, depending on the length of the gel. Subsequently, the RNA was visualized on a UV gel imager, and excess gel was cut away; leaving ∼0.75 cm of gel around the outer edges of samples lanes will improve transfer accuracy. The RNA was transferred with 3M NaCl pH 1 (with HCl) to an nitrocellulose membrane for 90 minutes at 25°C. Post transfer, the membrane was rinsed in 1x PBS and dried on Whatman Paper (GE Healthcare). Dried membranes were rehydrated in Intercept Protein-Free Blocking buffer (LI-COR Biosciences) for 30 minutes at room temperature. After the blocking, the membranes were stained using IRDye IR800 streptavidin for 30 minutes at 25°C. Excess Streptavidin-IR800 was washed from the membranes using three washes with 0.1% Tween-20 in 1x PBS for 3 minutes each at 25°C. The membranes were scanned on a LI-COR Odyssey CLx scanner (LI-COR Biosciences). Images and intensity of bands were acquired using LI-COR Image Studio software.

### sialoglycoRNA labeling and in vitro enzyme digestions

We isolated small RNA and labeled Ac_4_ManNAz with copper-free click using dibenzocyclooctyne-PEG4-biotin (DBCO-biotin, Sigma) as previously described^1^. After labeling RNA was analyzed via gel as described above. For in vitro heparinase digestions, 2 µg of small RNA was reacted in a final volume of 20 µL with 1x Heparinase Buffer (NEB) and 0.5 µL of each of the three heparinase enzymes described above. After 45 minutes at 37°C, the RNA was cleaned up with a Zymo column and analyzed by RNA blotting.

### Western blot

Cells were quickly rinsed with ice-cold PBS, and directly lysed with samples buffer (150 mM NaCl, 50 mM Tris, 0.5% TritonX-100, pH 7.4) containing phosphatase inhibitor cocktail (Cell Signaling Technology) on ice for 15 minutes. After centrifugation at 12,000x g for 15 minutes at 4°C, lysates were heated at 95°C for 10 minutes in 1x NuPAGE LDS loading buffer (ThermoFisher Scientific) containing 5 mM DTT. Samples were then resolved by SDS-PAGE using AnyKD Criterion TGX Precast Midi Protein Gels (Bio-Rad Laboratories) and transferred to nitrocellulose membranes. Membranes were blocked in blocking buffer, and incubated with primary antibodies (diluted in blocking buffer) at 4°C overnight. After washing three times for 3 minutes each in 1x PBS with 0.1% Tween-20 (PBST), membranes were incubated with secondary antibodies at room temperature for 45 minutes, followed by the same 3x PBST washing. Membranes were finally rinsed in 1x PBS and scanned on a LI-COR Odyssey CLx scanner. Images and intensity of bands were acquired using LI-COR Image Studio software.

Primary antibodies used: mouse monoclonal anti-EXT2 (Santa Cruz, sc-514092, immunoblot 1:1000), mouse monoclonal anti-GAPDH (Santa Cruz, sc-47724, immunoblot 1:1000), rabbit polyclonal anti-GFP (Invitrogen, A11122, immunoblot 1:1000), mouse monoclonal anti-NDST1 (Santa Cruz, sc100790, immunoblot 1:1000), mouse monoclonal anti-HS6ST1 (Santa Cruz, sc-398231, immunoblot 1:1000), mouse monoclonal anti-HS2ST1 (Santa Cruz, sc-376530, immunoblot 1:1000), rabbit monoclonal anti-phospho-p44/42 MAPK (Erk1/2) (Thr202/Tyr204) (Cell Signaling Technology, 4696S, immunoblot 1:500), mouse monoclonal anti-p44/42 MAPK (Erk1/2) (Cell Signaling Technology, 4370S, immunoblot 1:500). Secondary antibodies used: IRDye 800CW goat anti-Rabbit IgG secondary antibody (LI-COR Biosciences, 926032211, immunoblot 1:1000) and IRDye 800CW goat anti-Mouse IgG Secondary antibody (LI-COR Biosciences, 926-32210, immunoblot 1:1000).

### Genome-wide CRISPR/Cas9 screening

MOLM-13 cells expressing Cas9 under blasticidin selection^23^ were grown as above and selected with 10 µg/mL blasticidin for 3 days to ensure a homogenous starting population. Starting after selection on Day 0, 30 million cells were infected with 1:150 the genomewide-sgRNA lentivirus^23^ in 60 mL of fresh media with 8 µg/mL polybrene. On Day 3 cells were spun down and resuspended in 70 mL of fresh media with 1 µg/mL puromycin to select for sgRNA infected cells. On Day 5 the media was exchanged for fresh media with 1 µg/mL puromycin. On Day 6 the cells were switched to normal media without puromycin for expansion. From Days 7 to 18 the cells were counted and passaged as needed in fresh media to maintain a cell density between 750,000 and 2,000,000 cells per mL. On Day 18 cells were counted: 40M cells were saved prior to sorting for an input population reference and 200M cells for each Siglec-11 and 9D5 were saved for staining and sorting. To stain, 1000 µg Siglec-11 was precomplexed with 270 µg secondary antibody, 450 µg 9D5 was precomplexed with 225 µg secondary antibody, and 266 µg of MAAI was precomplexed with 266 µg Streptavidin AF647. Live cell staining was performed as noted above with Fc blocking; however before FACS sorting, cells were filtered over a 40 µm strainer (Corning) and stained with DAPI. Cells were selected for DAPI negative (live) and then the bottom 5% intensity of cells stained with each Siglec-11 or 9D5 were sorted into tubes. Cells were collected in FACS buffer after sorting spun down into pellets at 500x g for 4 minutes at room temperature; input cells were processed in a similar fashion to obtain a cell pellet. After removing the supernatant, cell pellets were frozen at -80°C for later processing. Cells were then processed by resuspending in 200 µL 1x PBS + 5 µL RNaseA + 20 µL Proteinase K and incubated at 25°C for 5 minutes. Then 200 µL of Buffer AL (Qiagen) was added and samples were carefully vortexed to mix without shearing genomic DNA (gDNA). The samples were then headed to 56°C for 60 minutes. After heating, 200 µL of 100% ethanol was added, gentle vortexing was again used, and then the material was purified over Zymo columns. All spins were performed at 6,000x g for 20 seconds: after spinning the sample through, the columns were washed twice with 80% ethanol and then the DNA was eluted with 2x 15 µL water. For input samples the initial volumes were scaled up from 200 µL to 1000 µL of digestion and Qiagen buffers.

To amplify the sgRNAs out of the gDNA we performed real-time PCR and for each sample performed 12 parallel reactions each with 1 µg of input gDNA. The PCR reactions were 50 µL final with 200 nM forward and reverse primers with 1x Q5 PCR Master Mix (NEB); 23 cycles of 98°C for 20 seconds, 65°C for 20 seconds, and 72°C for 90 seconds were completed. After PCR, the 12 reactions were pooled and purified over a Zymo column following the manufacturer’s recommended protocol for PCR DNA. To add a final index primer for sequencing, a final round of PCR was performed by taking 100 ng of amplified sgRNA library and 5 cycles of the above PCR program was run followed by Zymo column clean up. The finally indexed libraries were assessed for size and concentration on a BioAnalyzer High Sensitivity DNA Chip (Agilent). Libraries were pooled equimolar and then sequencing on the NextSeq platform (Illumina) with a 19 bp Read 1 and two 8 bp index reads. For the Read 1 a custom sequencing primer 5’- TCTTCCGATCTCTTGTGGAAAGGACGAAACACCG-3’ was used. Enrichment of guides and genes were analyzed using the MAGeCK statistical package^43^ by comparing read counts from each cell line with counts from matching plasmid as the initial population.

### Generation and characterization of KO and stably expressing cell lines

All CRISPR-Cas9 knockout assays used PX459^44^. The target oligonucleotides used were: EXT2: TCTCCCGGGAGTATAATGAA; NDST1: CCGGAGGCTGTGTCGGCACG; HS6ST1: CTACCTGAGCGAGTGGCGGC; HS2ST1: AATTGAGCAGCGACATACAA. U2OS cells were transfected with gDNA vectors. Two days later, puromycin (Invivogen, ant-pr-1) was added to the cell culture at a final concentration of 2 µg/mL and the live cells were selected by flow cytometry (BD science, FACS Calibur 2) for isolation of single clones. The expanded individual clones were screened by genomic DNA sequencing and western blot analysis.

Complementary DNA (cDNA) for human EXT2 was amplified from cDNA library (Takara Bio); cDNAs for Sulf1 and Sulf2 were gifts from Steven Rosen (Addgene plasmid)^34^. All three cDNA were inserted into mEmerald-C1 (Addgene). All plasmids were verified by DNA sequencing. To generate U2OS cells stably expressing mEmerald-EXT2, mEmerald-EXT2 D517N/D573N, mEmerald-Sulf1, or mEmerald-Sulf2, cells were transfected with the indicated plasmids and selected using 200-1,000 µg/µL (gradually increasing) G418 (Invivogen) for two weeks; green-positive cells were sorted into mono-clones by flow cytometry and cultured in the presence of 200 µg/µL G418 for 2 weeks. Proliferated clones were verified by immunoblotting and fluorescence imaging.

### UV crosslinking

25 ng/mL of VEGF-A_165_ were added to starved HUVECs for 5 minutes. Cells were treated by UV (60000 µJ, 2 minutes) on the ice and then directly lysed with samples buffer. For RNase treatment, RNaseA and RNaseIII were added to the samples at a final concentration of 1000 ng/mL and 20 U/mL, separately. Samples were all incubated at 37°C for 10 minutes and then lysed on ice for another 10 minutes. After centrifugation at 12,000x g for 15 minutes at 4°C, lysates were incubated with 5 µL Protein-G bead (Thermo Scientific) pre-conjugated with 1 µg of anti-VEGF-A (Proteintech) at 4°C overnight. The beads were washed three times with PBS and heated at 95°C for 10 minutes in 1x NuPAGE LDS loading buffer containing 5 mM DTT. Samples were then analyzed by Western blot described above. Anti-VEGF-A_165_ (R&D Systems, AF293-NA, immunoblot 1:1000) and donkey anti-Goat IgG secondary antibody (LI-COR Biosciences, 92632214, immunoblot 1:1000) were used as primary and secondary antibodies, separately.

### In vitro IP and rPAL

5 µL Protein-G bead was pro-conjugated with 1 µg of anti-VEGF-A or 9D5 antibodies in samples buffer for 1 hour at 4°C. After washing three times with samples buffer, beads were incubated with 1 µg of VEGF-A_165_ or VEGF-A_121_ for 2 hours at 4°C. The beads were washed three times with samples buffer and then incubated with 1 µg of HUVEC small RNA for 2 hours at 4°C. After washing three times with samples buffer, the beads were suspended in 50 µL RNA binding buffer and heated for 5 minutes at 50°C. Remove the beads and transfer the RNA extract solution to a new tube. Here, 100 µL of pure water was added and vortexed for 10 seconds, and then 300 µL of 100% ethanol was added and vortexed for 10 seconds. The RNAs were purified over a Zymo column.

For rPAL labeling, experiments were performed as described previously^18^. Briefly, lypophilized RNAs were suspended with 28 µL blocking buffer (1 µL 16 mM mPEG3-Ald (BroadPharm), 15 µL 1 M MgSO_4_ and 12 µL 1 M NH_4_OAc pH5 (with HCl)) and then incubated for 45 minutes at 37°C. 1 µL 30 mM aldehyde reactive probe (Cayman Chemicals, ARP/aminooxy biotin) is added first, then 2 µL mM NaIO_4_ (periodate) is added. The periodate is allowed to perform oxidation for exactly 10 minutes at room temperature in the dark. The periodate is then quenched by adding 3 µL of 22 mM sodium sulfite. The reaction is allowed to proceed for 5 minutes at 25°C, and then moved to 35°C for 90 minutes. The reaction is then cleaned up by Zymo column. The RNAs were eluted from the column using 2X 6.2 µL water and denatured at 55°C for 10 minutes with 12 µL of Gel loading Buffer II. Immediately after the heating, the RNAs were placed on ice for 2 minutes. Samples were then analyzed by RNA northern blotting and Streptavidin staining described above.

